# The functional role of oscillatory dynamics in neocortical circuits: a computational perspective

**DOI:** 10.1101/2022.11.29.518360

**Authors:** Felix Effenberger, Pedro Carvalho, Igor Dubinin, Wolf Singer

## Abstract

The dynamics of neuronal systems are characterized by hallmark features such as oscillations and synchrony. However, it has remained unclear whether these characteristics are epiphenomena or are exploited for computation. Due to the challenge of specifically interfering with oscillatory network dynamics in neuronal systems, we simulated recurrent networks (RNNs) of damped harmonic oscillators in which oscillatory activity is enforced in each node, a choice well-supported by experimental findings. When trained on standard pattern recognition tasks, these harmonic oscillator networks (HORNs) outperformed non-oscillatory architectures with respect to learning speed, noise tolerance, and parameter efficiency. HORNs also reproduced a substantial number of characteristic features of neuronal systems such as the cerebral cortex and the hippocampus. In trained HORNs, stimulus-induced interference patterns holistically represent the result of comparing sensory evidence with priors stored in recurrent connection weights, and learning-induced weight changes are compatible with Hebbian principles. Implementing additional features characteristic of natural networks, such as heterogeneous oscillation frequencies, inhomogeneous conduction delays, and network modularity, further enhanced HORN performance without requiring additional parameters. Taken together, our model allows us to give plausible a posteriori explanations for features of natural networks whose computational role has remained elusive. We conclude that neuronal systems are likely to exploit the unique dynamics of recurrent oscillators networks whose computational superiority critically depends on the oscillatory patterning of their nodal dynamics. Implementing the proposed computational principles in analog hardware is expected to enable the design of highly energy-efficient and self-adapting devices that could ideally complement existing digital technologies.

---

Neuronal networks of the cerebral cortex and the hippocampus and likely also those in homologue structures of non-mammalian species are characterized by several canonical anatomical and physiological features. Among those are recurrent connections between network nodes within and between processing layers [1, 2], the propensity of nodes and subnetworks to oscillate in different preferred frequency ranges [3, 4], heterogeneous but tuned conduction delays [5, 6], and activity-dependent adjustment of the gain of connections [7, 8] by Hebbian synaptic plasticity [9].

Certain canonical circuit motifs, such as recurrent inhibition and excitation, play a well-defined role in signal processing. Examples are contrast enhancement, gain control, dynamic range expansion, and competitive interactions. However, such microcircuits also tend to oscillate and, when several such microcircuits interact, additional complex dynamical phenomena emerge. In simultaneous recordings from multiple network nodes, these dynamics manifest themselves as frequency-varying oscillations [10, 11], transient synchronization or desynchronization of discharges [12], resonance [13], entrainment [14], phase shifts [15], and traveling waves [16, 17, 18]. Yet, the functional significance of many of these dynamical phenomena has remained unclear. Although the role of oscillating neurons and microcircuits in the generation of motor patterns is well established [19], it remains a matter of discussion whether oscillations support computations in the context of cognitive processes in the cerebral cortex and, if so, how [20, 21, 22, 23, 24, 25]. Moreover, determining whether these dynamical phenomena serve a functional role for computations is notoriously difficult in physiological experiments because strategies for the identification of causal relationships based on loss or gain of function interventions fall short in such complex and highly integrated systems [26]. Therefore, virtually all evidence for the functional role of the aforementioned dynamical phenomena has remained correlative in nature.

To overcome this epistemic hurdle and isolate computational principles, we performed in silico simulations of recurrent networks (RNNs) and trained them on standard pattern recognition tasks (sequential and permuted MNIST digit recognition and a time-series prediction task, see Methods and Supplementary Materials), which allowed us to use task performance as a measure for functional relevance.

Inspired by physiological evidence [27], we configured network nodes as damped harmonic oscillators (DHOs), a quintessential model of oscillatory dynamics that captures two fundamental properties of natural signals: harmonic oscillations and exponential decay. In the resulting Harmonic Oscillator Recurrent Networks (HORNs), few control parameters determine the relaxation dynamics of each network node. This choice acts as an inductive bias of nodal dynamics and permitted the comparison of networks in which nodal dynamics was non-oscillatory with various regimes of oscillatory nodal dynamics by solely adjusting the control parameters of the DHOs, leaving other network parameters unchanged.

Following this strategy allowed us to quantify the effect of oscillations on network performance. Furthermore, it helped bypass the issue that recurrent networks with non-oscillating nodes (e.g. leaky integrators) can exhibit oscillatory activity as an emergent property on the network level, complicating the analysis of the oscillations’ functional relevance per se.

We found that HORNs outperformed, sometimes by a large margin, their non-oscillating counterparts composed of leaky integrators, with respect to parameter efficiency, task performance, learning speed, and noise tolerance. This superiority was also manifest in comparison with other non-oscillating RNNs that rely on gated architectures (LSTM [28] and GRU [29]), and is particularly pronounced for small system sizes. The finding of increased task performance is supported by previous works, that also observed an increase in the performance of recurrent networks in the oscillatory regime [30, 31, 32, 33].

In-depth analyses of the dynamics of HORNs uncovered a powerful computational principle that uses the superposition and interference patterns of waves for stimulus representation and processing and that exploits the unique properties of coupled oscillator networks such as (de)synchronization, resonance, and nonlinear, frequency-dependent gain modulation, as well as the phase-based coding of stimulus sequence order. Without requiring fine-tuning of the parameters for the different experiments, the dynamics of HORNs shared numerous features with those observed in the cerebral cortex, suggesting that the uncovered computational principle is also exploited by natural networks. To further examine this possibility, we implemented other characteristic features of natural networks in HORNs and found that the inclusion of these biologically inspired features typically resulted in further improved task performance without increasing the number of trainable variables. Because the simulations allowed us to study the functional consequences of implementing known properties of natural neuronal networks – in particular of the cerebral cortex – our synthetic approach provided in addition plausible a posteriori explanations for a number of phenomena whose function has so far remained elusive or has given rise to controversial discussions.

In contrast to earlier research in neuroscience and machine learning, which focuses mainly on network dynamics and biological realism [34, 35, 36, 37] on the one hand, or task performance [31, 38, 39, 33] on the other, this study combines elements from both fields by simultaneously addressing the mechanistic modeling of biological network characteristics and their functional validation through benchmark tests.

## Results

### Oscillating network nodes

To permit analysis of the functional relevance of oscillatory dynamics in RNNs and to overcome the epistemic hurdle of emergent oscillations on the network level as described above, we enforced oscillations at each network node and configured the nodes of RNNs as damped harmonic oscillators (DHOs). Importantly, in contrast to traditional networks of non-oscillating nodes such as leaky integrators in which oscillations are merely an emergent phenomenon on the network level, the enforcing of oscillatory activity in every single node is quintessential for our approach, making the extended dynamical primitives available exclusively to oscillators (such as coding in phase space, (de)synchronization, and resonance, among others) stably available at each node. Choosing damped harmonic oscillators as the model of nodal activity was motivated by several reasons. First, DHOs represent the quintessential implementation of an oscillatory process in both physics [40] and neuroscience [41]. It furthermore allows full control of its relaxation dynamics through few, interpretable control parameters, making this model the fundamental choice for introducing oscillations at the level of a single network node. Second, damped harmonic oscillations in neuronal microcircuits generically result from excitatory-inhibitory interactions or negative feedback subject to a damping force [42, 43]. Third, recent experimental evidence showed that population activity in the visual cortex of macaques is well modeled by driven damped harmonic oscillators [27]. By manipulating the control parameters of the DHOs, we could convert the nodes from oscillators to integrators without interfering with other network properties, which allowed us to compare oscillatory with non-oscillatory regimes. Importantly, a DHO node in our networks should not necessarily be considered as a single biological neuron, but rather as a unit representing an abstract aggregate quantity of the activity of a microcircuit composed of recurrently coupled populations of excitatory and inhibitory neurons, such as a (P)ING circuit [44] or a cortical column [43]. Note that this abstract aggregate quantity could readily be considered as some form of local field potential, but also, for example, as an aggregate firing rate fluctuating around a base value, or as the fluctuation of the level of synchronous action potentials in a neural population. In this sense, HORNs should be understood as a mesoscale model of a cortical network (see Fig. 1A and Methods). Note that an alternative and functionally equivalent biological implementation of the oscillating nodes could have been single neurons endowed with pacemaker currents [45, 46, 47].

**Figure 1:**
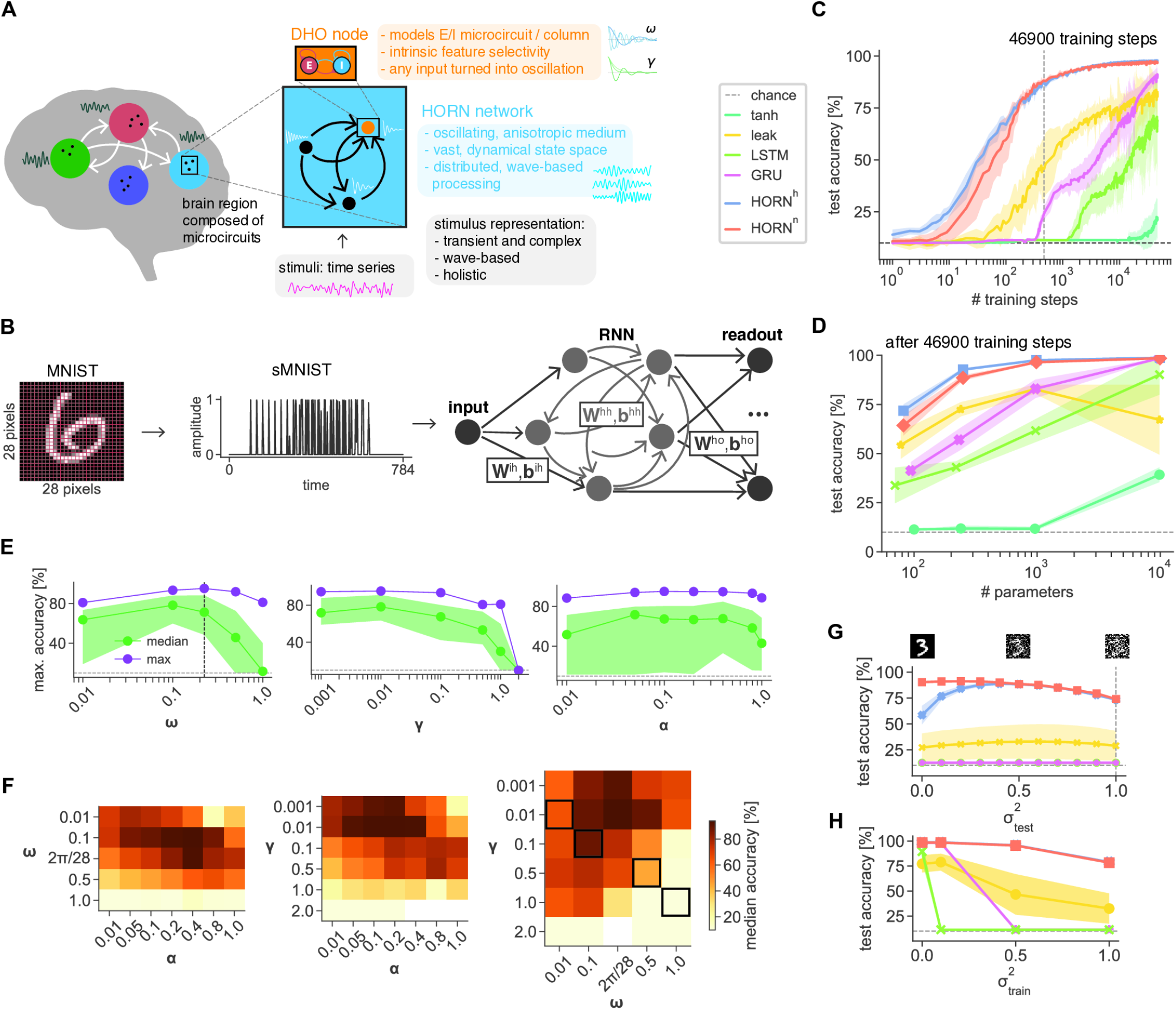
The architecture of HORN networks and their performance in the sMNIST pattern recognition task. **A**. A HORN network composed of DHO nodes as a mesoscale model of brain activity. Each DHO node models the compound activity of a recurrently connected E-I population as damped oscillations controlled by the parameters *ω, γ, α*. **B**. Serialization of an MNIST digit to an sMNIST stimulus and feeding the stimulus into an RNN with trainable parameters **W**^ih^, **b**^ih^ (input weights and biases), **W**^hh^, **b**^hh^ (recurrent weights and biases), and **W**^ho^, **b**^ho^ (readout weights and biases). For sMNIST, all networks are trained using a readout after 784 time steps. **C**. Test accuracy on sMNIST for different RNN architectures with approximately 2500 trainable parameters (tanh, leak, HORN 43 nodes, GRU 25 nodes, LSTM 22 nodes) as a function of training steps over 100 training epochs. Lines show mean accuracy over 10 network instances with random weight initialization, shaded areas standard deviation. Legend at the top left. Note the logarithmic scale in the number of training steps. **D**. Maximal test accuracy after 100 training epochs on sMNIST for different RNN architectures as a function of system size (number of nodes for approximately 10^4^ training parameters: tanh, leak, HORN 94 nodes, GRU 54 nodes, LSTM 47 nodes). Networks and legend as in C. Note the high task performance of HORNs at very low node and parameter counts. **E**. Distributions of maximum test accuracy on sMNIST of HORN^h^ networks (32 nodes, 10 training epochs) with variable parameter values *ω* (natural frequency), *γ* (damping factor), *α* (excitability). Each panel marginalizes the values for the remaining parameters. Lines show median and maximum, shaded areas indicate 25% and 75% percentiles. Note that maximal network performance peaks around a value of *ω* = 2*π/*28 (dashed vertical line, see main text). **F**. Median test accuracies of HORN^h^ networks as in E. Each matrix entry shows the median test accuracy (color scale on the right) for the marginalized remaining parameter. Parameter value pairs (*ω, γ*) resulting in critical damping (*ω* = *γ*) marked with a black frame. Note the performance drop for damping values above the critical point. **G**. Maximal test accuracy of different RNN architectures (10^4^ trainable parameters) after training on noisy sMNIST for 100 epochs as a function of training noise level 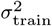. Lines show mean accuracy over 10 random networks, shaded areas show standard deviation. Colors as in C. **H**. Test accuracy of different network architectures trained on noisy sMNIST with 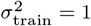 (dashed vertical line) as a function of test stimulus noise level 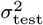, calculated over *n* = 1000 test stimuli. Colors as in C. Note the pronounced noise resiliency in HORNs.

In our model, each DHO node has one state variable *x*, the oscillator’s time-varying amplitude, and three control parameters that jointly determine its relaxation dynamics: the natural frequency *ω*, the damping coefficient *γ*, and an excitability coefficient *α* (see Methods). These parameters can be thought of as the innate or long-term adaptation of the node to the characteristics of the signals to be processed (Fig. 1A). Similarly as in works on neural ODEs and oscillator RNNs [48, 31, 49], a node is defined by the second-order ODE 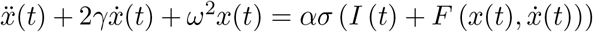, where *t* denotes time, *σ* is an input nonlinearity that models local resource constraints, *I*(*t*) denotes the time-varying input to the node, and 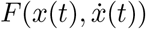 models feedback input of the node to itself that captures characteristics of the dynamics of the microcircuit that a DHO node represents (see Methods).

Despite their simplicity, DHO nodes capture the essential dynamical characteristics of various neural mass models in the oscillatory regime [50, 51, 42] (see Supplementary Materials). Thus, the oscillatory properties of the nodes implemented in our simulations are both physiologically plausible and experimentally confirmed [27].

It is important to note that already a single DHO node possesses intriguing computational capabilities that are not accessible to non-oscillating nodes such as leaky integrators. First of all, DHOs convert any input signal into an oscillation, even if the signal is non-oscillatory (this is the inductive bias on nodal dynamics in HORNs, see Discussion). Using an oscillatory representation enables a node to encode information not only in amplitude but also in phase, and in particular allows the encoding of stimulus sequence order in phase shifts (Fig. S1). If inputs are oscillatory, which is the case for recurrent inputs from other network nodes, but also for some sensory inputs, additional computational capacities can be leveraged. The DHOs then gain modulate the input signals in a nonlinear way as a function of their frequency (see Fig 2H, Fig. S2, S3 and Supplementary Information). Moreover, the interaction of the input nonlinearity with the biologically inspired feedback connections endows the DHO nodes with a dynamical repertoire that goes beyond that of a classical damped harmonic oscillator. For example, the feedback connections allow the nodes to express self-sustained oscillations and to resonate at fractional harmonics of their natural frequency (see Fig. S2, S3, S4 and Supplementary Materials).

**Figure 2:**
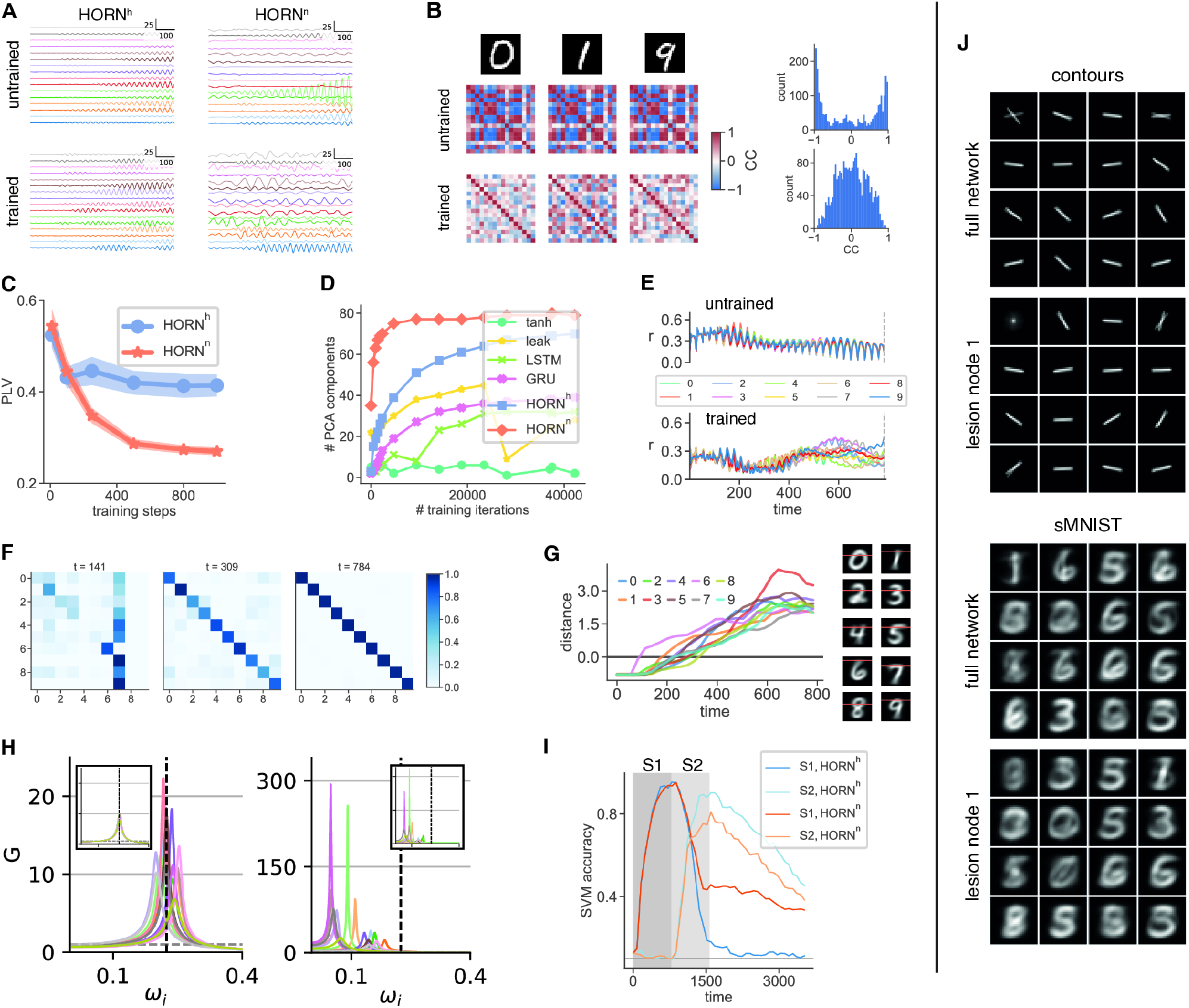
Dynamics and computational principles in HORNs. **A**. Network activity elicited by an sMNIST stimulus (digit 0) of 16-node HORNs trained on sMNIST, before (top row) and after (bottom row) training. Left column 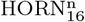, right column 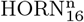. Note the highly irregular temporal dynamics of 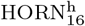 in the trained state. **B**. Pairwise cross-correlation coefficients of node activity for different input stimuli in 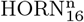 before (left, top row) and after training (left, bottom row) on sMNIST for 10 epochs. Distribution of cross-correlation coefficients before (right, top) and after (right, bottom) training. **C**. Pairwise phase locking value (PLV) calculated on *n* = 100 test stimuli as a function of training steps for a 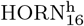 (blue) and 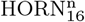 (orange) network trained on sMNIST. Lines show mean, shaded region indicates standard deviation. **D**. Number of PCA components explaining 99% of the variance of observed network activity as a function of training steps for different RNN architectures (10^4^ trainable parameters). PCA performed on network activity data resulting from 10000 test stimuli (see Methods). **E**. Dynamics of mean Kuramoto order parameter *r* of 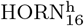 activity trained on sMNIST in the untrained network (top) and after training for 10 epochs (bottom). Lines show mean values per digit class. The HORN learns to generate dynamic, stimulus-specific patterns of higher-order synchrony. **F**. Normalized confusion matrices of linear SVMs trained on stimulus classification from activity of a 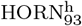 (trained on sMNIST for 100 epochs) activity at different time points of the stimulus presentation period. Matrices show relative frequencies the number of predictions obtained over 10^4^ samples. The vanishing of the non-diagonal entries (indicating misclassified stimuli) shows that the network performs evidence accumulation over the stimulus presentation time. **G**. Mean distance to SVM decision boundary for SVMs as a function of presentation time for the dataset depicted in F. The color of the curves refer to different digits (insert). Note the steady accumulation of evidence. Right: Class-averaged MNIST digits with the image row corresponding to the decision boundary crossing of the SVM colored in orange. **H**. Intrinsic RFs (gain functions *G*) of nodes as a function of input frequency *ω*_*i*_ of 16-node HORNs before (insets) and after training on sMNIST. Homogeneous (left) and heterogeneous (right) HORN variants. Gain values *G >* 1 indicate amplification. Horizontal dashed lines indicate *G* = 1, vertical dashed lines indicate the frequency *ω*_*v*_ = 2*π/*28 that corresponds to a straight vertical bar in pixel space. Colors indicate node identity. **I**. Accuracy of stimulus decoding by linear SVMs trained on the activities of different HORN networks (93 nodes) that were stimulated with a sequence of two temporally segregated stimuli S1, S2 (shaded areas indicate stimulus timing). Light and dark colored curves indicate SVM decoder performance for S1 and S3 for a homogeneous (HORN^h^) and non-homogeneous (HORN^n^) network, respectively. Note the longer persistence and superposition of stimulus-specific information in the HORN^n^. **J**. Effective RFs of a 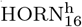 when mapped with contour stimuli from LSDSa (top two panels) and sMNIST digits (bottom two panels). Both cases show the ERFs of the full network and the ERFs after the deletion (lesion) of the first node. Note the reconfiguration of most of the ERFs when a node is inactivated.

The natural frequency parameter *ω* defines a frequency band in which the DHO node selects and amplifies temporally modulated signals through resonance (see “Receptive Fields” below). When integrated in a recurrent network, DHOs convert inputs into wave patterns. These patterns manifest both as standing waves at each network node and also propagate throughout the whole network, allowing for interference with waves generated by spatially and temporally segregated stimuli, as well as those generated by internal and reverberating network dynamics. Note that although networks consisting of leaky integrators can also support propagating traveling waves, in this case interference is limited to the transient interactions between wave fronts, as nodes cannot represent standing waves in the way that DHOs can. Crucially, when input signals exhibit distinct temporal patterns, the oscillatory behavior of DHO units enables them to identify features via resonance, rather than merely through integration over converging input connections. The latter strategy is commonly used in non-oscillating RNNs and all feed-forward architectures [52]. In HORNs, single nodes or assemblies of nodes can further encode information in distinct frequency bands and gate the flow of information as a function of relations among frequencies. These features of DHOs endow HORNs with the properties of an anisotropically coupled, nonlinear, analog medium for information processing that can exploit for its computations wave-based representations of stimuli and their interference patterns, which allows for a rich and high-dimensional coding space.

The choice of modeling neural population activity on a mesoscale with a DHO node allows us to uncover the generic principles described in this work. However, this approach limits the model’s ability to capture certain dynamical phenomena observed in neuronal systems. Among these are dynamical variations in oscillation frequency, spikes and their bursting behavior, and more generally any dynamics which are non-oscillatory or cannot be captured at the population level [53, 50]. Still, some of those dynamics could be modeled if the parameters *ω, γ, α* that influence the relaxation dynamics at each node are allowed to be driven by the dynamics of the network itself.

### Configuration and training of the networks

We first consider homogeneous HORN (HORN^h^) networks with all-to-all connectivity, no conduction delays, and identical parameter values *ω, γ, α*, for all nodes. As cortical circuits operate in a balanced state [54] where excitatory and inhibitory drives cancel in the mean (E/I balance), emphasizing the significance of fluctuations, we chose to couple DHO units on their velocity term 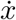 (see Methods). Stimuli were presented to the networks in the form of time series. Due to its standard nature and widespread use, we chose the MNIST handwritten digit classification task as our default benchmark task. We have also trained HORN networks on more challenging datasets such as a spoken-digit classification task, confirming the computational superiority of the oscillatory dynamics implemented in HORNs over non-oscillating architectures also in those cases; see Supplementary Materials. To transform an MNIST image into a time series, henceforth addressed as a sequential MNIST (sMNIST) stimulus, intensity values were collected in scan-line order from top left to bottom right of the images, turning a 28×28 pixel MNIST digit into a time series of length 784 (see Fig. 1B and Methods). This transformation converts geometrical properties of the stimuli into spectral patterns that can be exploited by HORNs to perform stimulus classification. Note that this approach serves as a proxy for other spectral regularities typical for natural signals and that we also study the case of geometrically organized inputs in the following (see “Geometric input drives self-organization”). To increase task difficulty, we also trained networks on permuted sequential MNIST (psMNIST) stimuli in which a random, but fixed permutation is applied to the time series. This operation destroys local luminance correlations in MNIST digits and results in time series stimuli with a flat spectrum, making frequency-based information processing harder (Fig. S5).

The networks are then trained on the 10-class stimulus classification task in a supervised manner, using an affine readout at the last time step trained with the back propagation through time (BPTT) algorithm (see Methods). For sMNIST, a grid search was performed on the triplet of parameter values *ω, γ, α* to find a constellation of DHO control parameter values that resulted in high classification performance (Fig. 1E,F, Table S1, and Methods).

The optimal parameter values turned out to be insensitive to network size (data not shown) and networks with low values of the damping factor *γ* tended to perform best (Fig. 1E, F). Low damping factors put the system in a highly oscillatory state and enhance the memory spans of individual DHOs as well as of the whole network (Fig. 1E,F), see also [55]. The excitability parameter *α* had no strong effect on performance, with intermediate values between 0.2 and 0.4 resulting in the best performance (Fig. 1E,F). The optimal values of the natural frequency parameter *ω* were found to be around the value *ω*_*v*_ = 2*π/*28 that corresponds to the fundamental frequency of an sMNIST time series resulting from a straight vertical line in the 28×28 MNIST pixel space (see Fig. 1E,F and Methods). As DHOs resonate with input frequencies around *ω*_*v*_, this choice of *ω* enabled the nodes to extract and retain the information contained in time series corresponding to continuous lines of different slopes in pixel space. The value *ω*_*v*_ also coincides with a peak in the variance of power spectral densities (PSDs) calculated on a representative set of sMNIST samples (Fig. S5). Thus, by adjusting *ω*, the network can be adapted to the statistical regularities of the sMNIST stimuli and this setting of priors enhances performance by allowing feature extraction through resonance rather than through selective recombination of convergent input connections. To the best of our knowledge, this feature extraction strategy has not been previously proposed and crucially depends on the presence of units or microcircuits with a propensity to oscillate.

### Network performance

Endowing network nodes with oscillatory dynamics improved network performance compared to leaky integrator or gated unit networks with respect to learning speed, parameter efficiency, and noise tolerance (Fig. 1C,D). Furthermore, we found that HORNs also outperform state-of-the-art gated RNN architectures in learning speed (Fig. 1C, Fig. S6), performance, and noise tolerance, at the same number of learnable parameters. This superiority was particularly pronounced in the region of low parameter counts (see Fig. 1D) and critically depended on the configuration of the network nodes to be in a highly oscillatory state (*ω* ≫ *γ*, see Fig. 1F, rightmost panel). Performance was found to drop for networks that were given very low oscillation frequencies (*ω* ≪ 1), which act more like (leaky) integrators, and for damping values above the critical value *γ > ω* that abolish nodal oscillations (see Methods). This shows the functional benefit of the presence of oscillating nodal dynamics in HORNs. When making the task more difficult by decreasing signal-to-noise ratios or increasing the number and similarity of stimulus patterns, the performance differences between HORNs and the other networks tended to increase further (Fig. S6, S7, S8, S9).

Note that even a simple leaky integrator network showed higher convergence speeds than an Elman RNN and the gated architectures when hyperparameter values were optimized by a grid search, albeit with a lower final test accuracy and with a much higher sensitivity to the random weight initialization. This finding can be explained by the fact that both HORNs and leaky integrator networks benefit from the residual connections introduced by the discretization scheme of the underlying ODE. This scheme introduces stable Lyapunov exponents and stabilizes gradients [55]. Choosing sufficiently small values of *ω* and *γ* results in stable gradients, which facilitates training by BPTT (see Methods). The oscillatory activity in HORNs also allowed the nodes to extract features by means of resonance and acted as an inductive bias. This bias allowed the network to exploit information present in the spectrum of sMNIST digits when the intrinsic nodal frequencies were tuned to the dataset (see “Receptive Fields”, Methods, and Fig. S11). A more detailed discussion of how the dynamical primitives available to HORNs lead to the observed pronounced increase in task performance is given in the next sections.

In contrast to the other architectures tested, the HORNs were noise resistant and showed only a gradual decline in task performance with increasing noise levels (Fig. 1G). Furthermore, the stimulus representations in HORNs were robust to a mismatch of noise characteristics between training and inference runs (Fig. 1H). This robustness can be explained by the strong attenuation of high-frequency signals due to the nonlinear, frequencydependent gain modulation of input signals at each DHO node (see “Receptive Fields”). In particular, this noise tolerance persists even when HORNs are trained on shuffled MNIST stimuli (psMNIST), in which the dominance of the low-frequency information characteristic for the sMNIST stimuli is removed (Fig. S10, S5). As both biological and artificial systems have to be able to learn from noisy stimuli and robustly detect, classify, and process stimuli even under changing noise characteristics, this noise resilience constitutes another interesting property of HORNs. To assess the influence of the presence of feedback parameters *v, w* on network performance, we trained HORNs with and without DHO feedback connections on sMNIST and psMNIST and measured their task performance (Fig. S12, S13). We found that even in the absence of feedback in both amplitude and velocity (*v* = *w* = 0), HORN performance decreased only slightly, but both feedback terms were needed to obtain the fastest learning speed and the highest overall task performance.

As expected, before learning, the dynamics of homogeneous HORN^h^ networks is dominated by large-scale synchronization among the nodes (Fig. 2A,B,C). As learning progresses, global synchronization decreases (Fig. 2C), increasing the dimensionality of network dynamics (Fig. 2D). This reduction in global synchronization is accompanied by the emergence of complex, spatiotemporally structured correlations and higher-order synchronization patterns that are stimulus specific, well segregated in the high-dimensional activity landscape of the network, and classifiable by a linear read-out (Fig. 2F,G).

### Heterogeneous networks

The structural and functional organization of mature cortical networks is characterized by heterogeneity [56, 57]. Although several recent studies found that this variability can be beneficial for learning and computations [58, 59, 60, 61], it is still debated to what extent natural heterogeneity is functionally relevant. To test whether increasing network heterogeneity facilitates learning in HORNs, we simulated non-homogeneous HORNs (HORN^n^) in which each node had a different natural frequency, damping coefficient, and excitability (see Methods). As expected, heterogeneous HORNs responded already in the untrained state with more complex and less globally synchronized patterns (Fig. 2A). As in the homogeneous case, global synchrony decreased further as learning progressed (Fig. 2C). Here again, the decrease in global synchrony led to an increase in the dimensionality of the dynamics (Fig. 2D), although the dimensionality of the dynamics was higher compared to their homogeneous counterparts already before training.

A comparison between homogeneous and heterogeneous HORNs revealed superior performance of the latter with respect to learning speed in data sets with more complex spectral structure such as psMNIST (Fig. S7). In addition, noise tolerance was also enhanced (Fig. 1G,H, Fig. S7). For such datasets, heterogeneous HORNs performed better than their parameter-optimized homogeneous counterparts. Note that in the case of sMNIST digits that have a reduced complexity of their signal statistics, the final performance of heterogeneous HORNs was found to be on par with that obtained by homogeneous HORNs whose parameters had been optimized for specific stimuli (Fig. 1C,D, S6). Thus, heterogeneity allows one to obtain networks with high task performance without the need to find optimal parameter configurations, saving computationally expensive resources. The higher task performance of heterogeneous over homogeneous networks results from the fact that heterogeneity introduces a multitude of timescales into network dynamics that can be used to process and represent stimuli [55]. Furthermore, heterogeneity brings the dynamics of the network closer to criticality [62, 63, 60, 55]. Network dynamics close to the critical point is characterized by scale invariance and divergence of spatial and temporal correlation lengths, leading to an increased dynamic repertoire and longer memory timescales. States close to criticality bear computational advantages for reservoirs that code information in transient states [62]. As HORNs also encode information in transients, the long-lived transient states enabled by dynamics close to criticality increase their computational power and allow them to best meet the trade-off between the efficiency and the stability of learning [55].

Generally, the advantages of heterogeneity were found to be especially prominent for larger networks, more difficult classification problems, and higher levels of stimulus noise (Fig. S6, S7, S8, S9). Because heterogeneous HORNs produced highly structured response landscapes already in the untrained state, we hypothesized that they might also serve as efficient reservoirs [64]. We found that, in contrast to RNNs composed of gated units that tend to have chance level stimulus decodability by a linear readout in the untrained state, heterogeneous HORNs already allow for high levels of decodability even in an untrained state (Fig. S14). To determine the reservoir properties of HORNs, we simulated heterogeneous HORNs with random but fixed recurrent weights and trained only the input and readout weights with BPTT. Although these reservoir HORNs showed a performance drop compared to the corresponding regular HORNs in which the recurrent connections were plastic (Fig. S15), we hypothesize that heterogeneous HORNs could serve as efficient reservoirs, especially with a larger number of nodes.

### Conduction delays

Another source of variability in biological neuronal networks are scattered conduction delays between nodes. In the cerebral cortex, neurons interact through relatively slow conducting nerve fibers (0.5 to 10 m/s), which introduces considerable, widely scattered, distance-dependent coupling delays [65]. To test the influence of introducing coupling delays on task performance, we started with a HORN^h^ and endowed all recurrent connections with uniformly distributed variable coupling delays [1,d_max_] (see Methods). This manipulation increased HORN performance in both maximal classification accuracy and learning speed on psMNIST (Fig. 3A). Increasing d_max_, which results in greater heterogeneity, was found to increase task performance, and this gain of function increased with increasing values of d_max_ (Fig. 3A). Thus, like for the preferred oscillation frequencies, heterogeneity in conduction delays enables the generation of more diverse spatiotemporally structured activity landscapes in HORNs, thus increasing the dimensionality of the networks’ state space. This again results in enhanced HORN performance, in particular for datasets with complex spectral properties such as psMNIST or changing noise characteristics (Fig. 1G,H).

**Figure 3:**
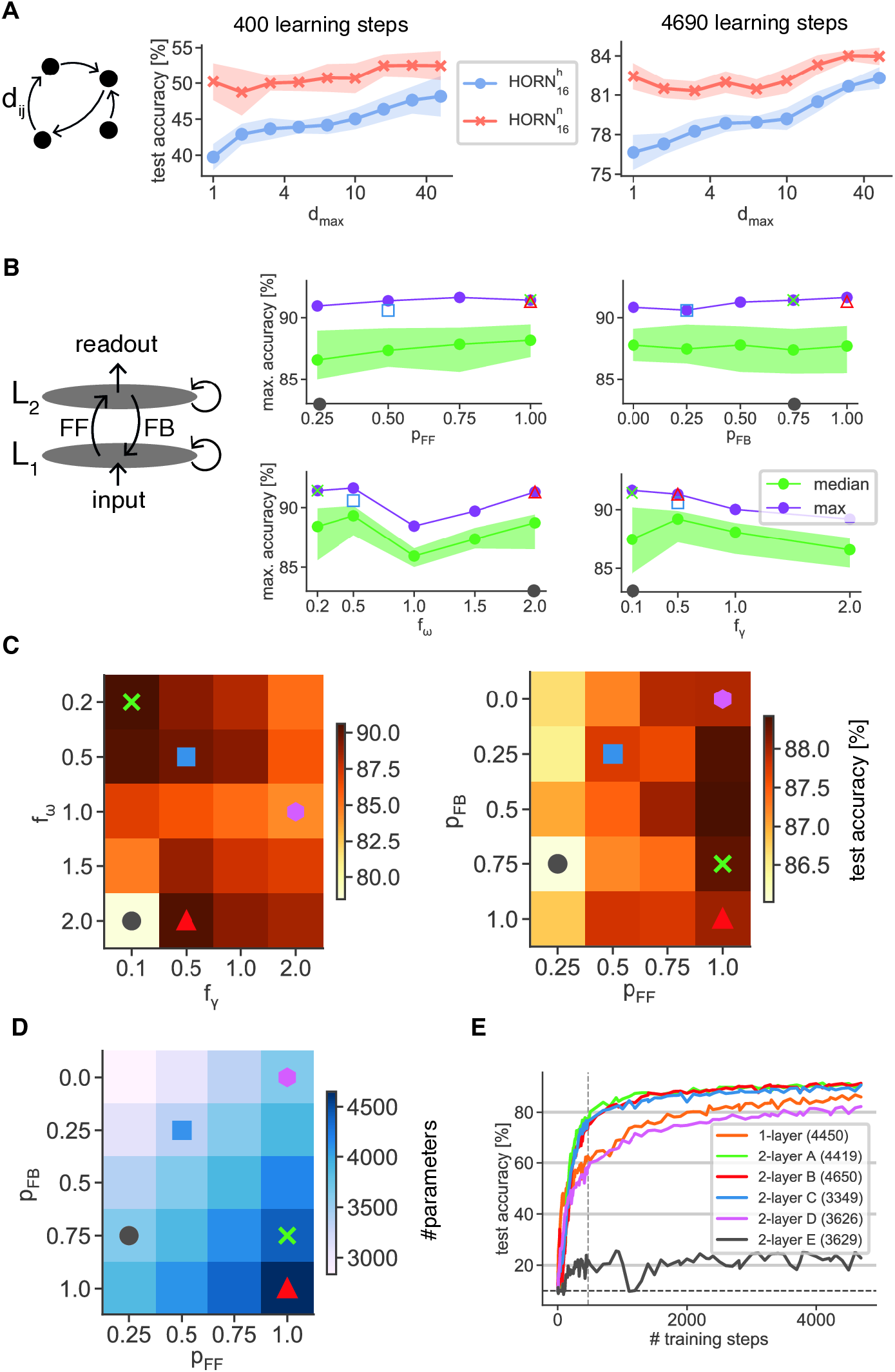
Biologically-inspired extensions to HORN networks. **A**. Performance of HORNs with connection delays. Maximum test accuracy on psMNIST after 400 training steps (left) and after 4690 training steps (right), respectively, as a function of maximal synaptic delay *d*_max_. For each network, connection delays were sampled from a uniform distribution *U* ([1, *d*_max_]). A network with *d*_max_ = 1 corresponds to a regular HORN. Lines show mean performance over 10 randomly initialized networks, shaded areas show standard deviation. **B**. Two-layer 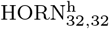 performance as a function of the 4 parameters *p*_FF_, *p*_FB_ (fraction of feed-forward and feed-back connections present, upper panels), *f*_*ω*_, *f*_*γ*_ (scaling factor of *ω* and *γ* between the first layer and the second layer, lower panels). Curves represent median and maximal values of marginal distributions of the maximal accuracy on psMNIST attained during training for each value of the 4 parameters. Colored symbols refer to five exemplary networks whose parameter constellations are shown in C. Note the drop in performance when L1 and L2 have the same preferred oscillation frequency and that increasing connection sparsity does not result in a strong performance drop (gray circle marked on axes lies outside of the accuracy range displayed). **C**. Median HORN^h^ performance (color scale on the right) for the different pairs *f*_*ω*_, *f*_*γ*_ values (left) and *p*_FF_, *p*_FB_ (right). Symbols refer to the same networks as in B. Note that different parameter combinations support equally high performance. **D**. Number of trainable parameters of HORN networks (color code on the right) as a function of various combinations of *p*_FF_, *p*_FB_. Symbols refer to the same networks highlighted in B and C. Note that certain network configurations show high performance despite lower numbers of trainable parameters (green cross, red triangle, blue circle). **E**. Evolution of accuracy as a function of learning steps of the 5 exemplar two-layer networks marked by symbols in B, C, D (colors of curves correspond to symbol colors) in comparison to a single layer HORN^h^ with a comparable number of trainable parameters. Number of trainable parameters in parentheses.

### Multi-layer networks

In the mammalian cerebral cortex, sensory signals are processed in hierarchically organized cortical areas that are reciprocally coupled [66]. To investigate the potential benefits of distributed multistage processing, we generate two-layer HORNs consisting of a “lower” (L1) and an “upper layer” (L2), with each layer consisting of a homogeneous HORN of 32 nodes, 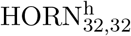, and introduced sparse reciprocal connections between the layers (Fig. 3B). The input signals were presented to L1 as before, and the result was read out at the nodes of L2. We performed a grid search on the inter-layer feed-forward and feed-back connection probabilities *f*_F_, *f*_B_, as well as the scaling factors *f*_*ω*_, *f*_*γ*_ that controlled the scaling of the parameters *ω, γ* of L2 with respect to L1. For each parameter configuration, we trained a 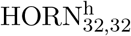 and evaluated task performance using the maximal test accuracy achieved on psMNIST in 10 training epochs (we chose psMNIST over sMNIST due to the much richer spectral stimulus properties). Note that for each fixed value of each parameter, there usually exists a parameter configuration that results in a high-performing two-layer network (Fig. 3B).

Importantly, we found that many two-layer 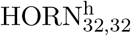 networks outperformed a single-layer HORN^h^ at a comparable number of trainable parameters (see Fig. 3C-E). In particular, higher task performance resulted when both the preferred oscillation frequency and the damping coefficient were lower in the upper layer (Fig. 3B, bottom row). In this setting, the dynamics of the faster lower layer L1 is partly opaque to the slower upper layer L2 due to the stronger attenuation of high frequencies in L2. However, activity in L2 is capable of entraining the nodes in L1, potentially supporting processes such as feature binding by contributing more global binding criteria available to the upper layer, but not to the lower layer. Because the upper layer nodes receive convergent input from the lower layer and operate at a longer time scale, the upper layers can bind longer segments of the stimulus time series.

Interestingly, we found that a two-layer network in which the separation of frequency bands across layers breaks down, fails to learn (Fig. 3E). In this case, the computations in the different layers are not sufficiently separated into frequency bands, and the cross-talk between layers not separated into different frequency bands hinders the networks to successfully solve the stimulus classification problem. For this reason, we consider it unlikely that multilayer recurrent networks without oscillating nodes can leverage these particular advantages of hierarchical processing.

### Fading memory and evidence accumulation

Due to the reverberation of activity, recurrent networks exhibit fading memory, which allows them to integrate information about successive events. Electrophysiological recordings of neurons in the visual cortex have shown that population activity exhibits fading memory and is capable of simultaneously representing classifiable information on the identity of successively presented stimuli, including their sequence order [67]. HORNs share this ability. Following the sequential presentation of two different stimuli, a linear classifier (support vector machine, SVM) can decode the two stimuli from the same segment of reverberating activity (Fig. 2I). As heterogeneous HORNs possess more diverse memory timescales than homogeneous networks, their ability to represent simultaneously information about temporally segregated stimuli is superior, yet another functional advantage of network heterogeneity (Fig. 2I). Further studies are required to determine whether non-oscillatory RNNs cope with the encoding of sequence order as effectively as HORNs.

As is characteristic of RNNs, the response patterns of HORNs evolve over time as a result of network dynamics. To determine the time point at which the network converges to states of maximal stimulus specificity, a linear SVM was trained on stimulus classification using activity data from a trained homogeneous HORN at different time points throughout the stimulus presentation period. The network accumulates evidence, with its dynamics allowing progressively better decoding of stimulus identity as the network approaches the read-out time *t* = 784 on which it was trained (Fig. 2F,G, S14).

### Receptive fields

To better understand the principles of computation in HORNs and how stimulus-specific activity patterns emerge during training, we investigated how learning changed the response properties of both individual nodes and the entire network. We expected analogies to the experience-or learning-dependent modifications of receptive fields (RFs) in neuronal systems [68]. Each DHO node in a HORN has a gain curve *G*(*ω*_*i*_) that describes how the node modulates the amplitude of a temporally modulated input signal as a function of the input signal frequency *ω*_*i*_ (Fig. 2H). Note that this holds both for external, stimulus dependent signals if they have temporal structure and always for the oscillatory activity conveyed by the intrinsic recurrent connections. The shape of G is determined by the values of *ω, γ*, and the self-connectivity terms *v* (the amplitude feedback) and *w* (the velocity feedback, given by the diagonal terms of the recurrent weight matrix) (see Methods, Fig. S3 and Supplementary Information). We call *G* the intrinsic receptive field (IRF) of the node because it defines the frequency band in which the node shows feature selectivity. During learning, the adjustment of the self-connection weights *v* and *w* alters *G* and therefore drives changes in the IRF of each node, allowing the node to improve its selectivity for stimulus features useful for performing a given task (Fig. 2H). In homogeneous HORNs where the values of *ω, γ, α* are the same for all nodes, the IRFs of all nodes only differ due to learning-induced changes in the values of the feedback parameters *v* and *w* that vary between nodes (Fig. 2H, left). For heterogeneous networks, the IRFs of the untrained network already cover a larger portion of the frequency space, with nodes tuned to a wider variety of features in the frequency space (Fig. 2H, right). Note that the value of *ω* not only influences the IRF but also sets the frequency band in which the node codes. This increases the diversity of frequency bands available for processing in heterogeneous HORNs. For comparison with neuronal systems, we mapped the receptive fields of the nodes in the same way as is common practice in electrophysiological studies [69]. We examined which stimuli activate a node most strongly and address the so-defined RF as effective RF (ERF). Like in neuronal systems [68, 70], the structure of ERFs varied greatly depending on the nature of the test stimuli. The reason is that responses depend not only on the external input but also on the network’s recurrent dynamics. We determined the ERFs of nodes in our models with two canonical choices of mapping stimuli: (i) simple line segments of different orientations, as commonly used for RF mapping in visual experiments [69] (Fig. S16), and (ii) the original MNIST digits on which the networks were trained, as an example of complex natural stimuli [70].

To measure ERFs, we simulated a heterogeneous 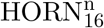 trained on sMNIST and calculated for each node the mean stimulus that resulted in maximal node activation (see Methods). When stimulated with simple line stimuli, we find simple orientation-selective ERFs that closely resemble the RFs of neurons in the primary visual cortex (Fig. 2J, top two panels). When mapped with complex stimuli, some nodes exhibited specificity for particular features of the MNIST digits (Fig. 2J, bottom two panels). Other nodes had ERFs that appeared to be unrelated to the stimuli in the training set, and yet others were inhibited. To determine the influence of the collective network dynamics on individual ERFs, we silenced one node of the network. This resulted in an immediate reconfiguration of the ERFs of the other nodes, which was quite dramatic in some cases (Fig. 2J). This indicates that after learning, ERFs are to a large extent the result of dynamic interactions in the network. Thus, the ERFs of HORN nodes undergo context-dependent dynamic modifications similar to those observed in natural systems [71].

To obtain a more quantitative description of the ERFs of the nodes in frequency space, we additionally stimulated the networks with harmonic sine-wave stimuli and determined the frequency responses of the nodes (Fig. S17). As expected, in untrained networks, most of the nodes resonated best with inputs whose frequencies were close to their preset natural frequencies. However, this preference changed substantially over the course of learning, with the nodes developing complicated and hard-to-predict resonance patterns, confirming the strong influence of anisotropic network interactions on the structure of ERFs.

### Learning priors

In the primary sensory cortices of the mammalian brain, information about statistical regularities of the natural environment is stored in the architecture and the synaptic weight distribution of both feedforward and reciprocal connections [2]. This translation of statistical contingencies of features into functional architectures is achieved during early development by experience-dependent synaptic pruning and continues throughout life through perceptual learning [2, 72]. The shaping of feedforward connections leads to canonical feature-selective RFs of the nodes. The selective stabilization of the recurrent connections among nodes leads to preferential coupling of nodes (representing, for example, cortical columns) tuned to features that frequently co-occur in natural environments. Hence, the weight configurations of the recurrent connections capture the statistical regularities of the natural environment, harboring the Gestalt criteria used for feature binding and perceptual grouping (see [2] for a review). To test how the installation of priors in HORNs influences their learning and task performance, we first installed a set of canonical priors by training a heterogeneous HORN^n^ (93 nodes) to discriminate simple elongated contours of different orientations placed at random locations in the MNIST pixel matrix (see Fig. S16 and Methods). We then froze the weights of the input and recurrent connections and trained only the readout layer on the sMNIST classification task with BPTT. We found that the installation of canonical priors – oriented lines – allowed the pretrained HORN to generate well separable representations of the MNIST digits. These representations were directly classifiable with high stimulus specificity by training only the readout layer (Fig. S18). When, after the installation of the priors, we only froze the weights of the input connections but allowed the prestructured recurrent connections to participate in further learning, we observed a further increase in learning speed (Fig. S18). In summary, these prestructured HORNs required substantially fewer training steps than HORNs trained directly with the MNIST digits to reach the same performance levels. This indicates that a priori knowledge of the statistical contingency of features substantially facilitates and accelerates learning classifiable object representations in HORNs. We noticed that pretrained heterogeneous networks required fewer training steps to reach high performance as their size increased (Fig. S7, S8). Therefore, we predict that sufficiently large and heterogeneous pretrained HORNs should be capable of single-shot learning, which is a hallmark of natural neuronal systems that have much higher complexity.

Taken together, these findings emphasize the beneficial effects that priors have for the orthogonalization of object representations in artificial and most likely also natural neuronal networks.

### Hebbian learning

To test whether gradient-based training of HORNs produces weight distributions compatible with those predicted by Hebbian learning principles (see above), we compared recurrent weight distributions and the corresponding correlation structure of responses before and after learning. To our surprise, we found that the changes in synaptic weights of HORNs induced by BPTT are similar to those predicted by a Hebbian mechanism (Fig. 4A,C). The BPTT learning algorithm apparently capitalizes on the stimulus-specific correlation structure of network activity and enhances those connections that induce correlation patterns characteristic of a particular stimulus. As expected, heterogeneous HORNs could exploit stimulus-specific correlation structures right from the beginning of training (Fig. 4C). Homogeneous HORNs, in contrast, must first learn to desynchronize to increase the dimensionality of their state space and to install stimulus specific correlation structures (Fig. 2A,B).

**Figure 4:**
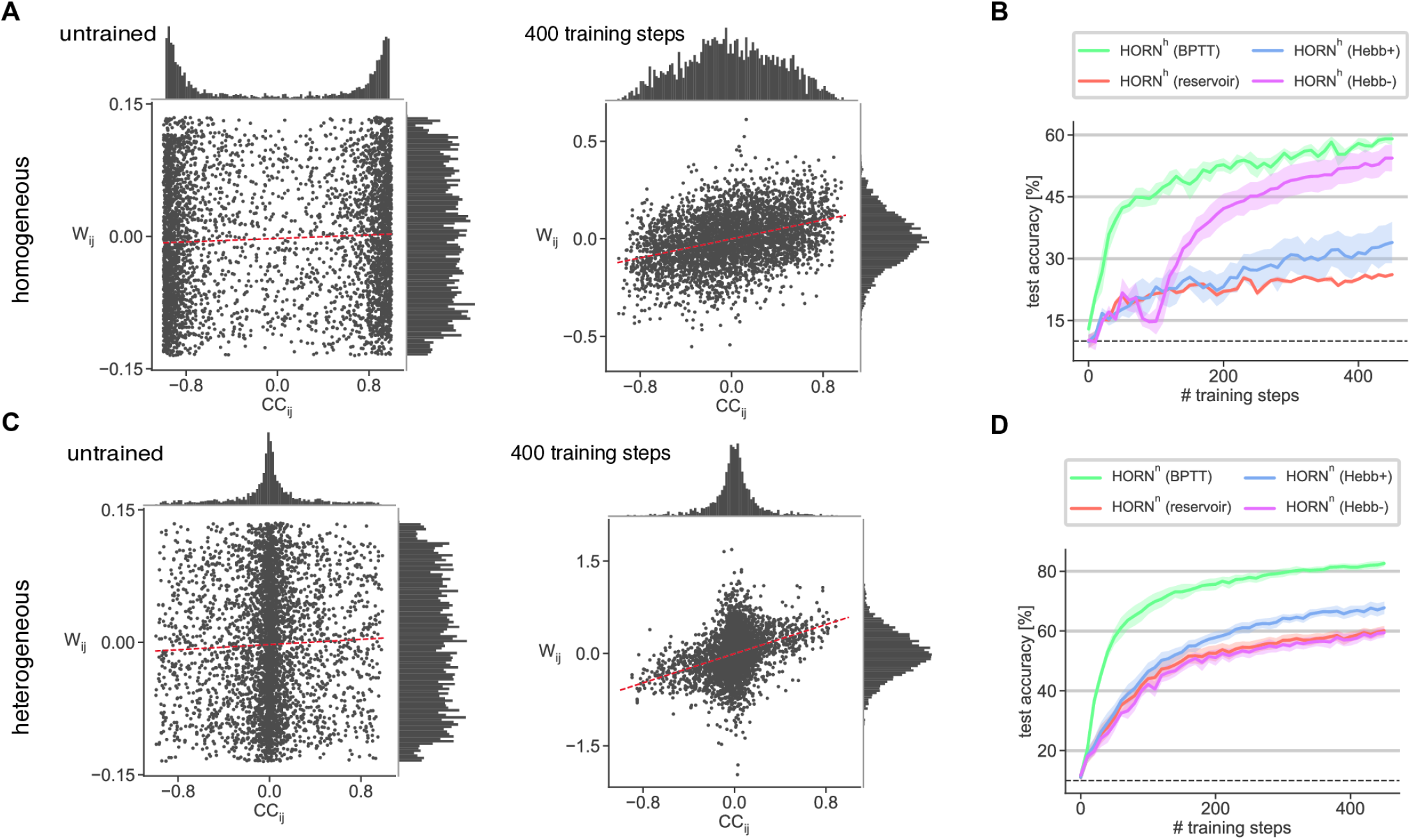
Backpropagation learning in HORN networks results in Hebbian-like weight changes. **A**. Scatter plots of connection weights 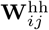 and mean cross-correlation coefficients CC_*ij*_ of node activities of a homogeneous HORN^h^ (64 nodes) before training (left) and after training on psMNIST for 500 training steps (right). CC_*ij*_ computed over 100 samples. Linear regression lines in red. Marginal distributions of 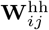 and CC_*ij*_ are shown on the right and top, respectively. Note the bimodal distribution of correlation coefficients with modes around -1, 1 in the untrained state and the more decorrelated network activity as a result of learning. **B**. Performance of a homogeneous HORN^h^ (64 nodes) network as a function of training steps when instances of the same network are trained with correlation-based Hebbian (suffix “Hebb+”) or anti-Hebbian (suffix “Hebb-”) learning rules compared to instances trained with BPTT (suffix “BPTT”) and when **W**^*hh*^ was fixed (suffix “reservoir”). The input and readout parameters are trained with BPTT for all instances. Curves show mean performance over 10 network instances with random weight initialization, shaded areas standard deviation. Note the strong performance of the anti-Hebbian rule for this initially highly synchronized network. **C**. Scatter plots of the connection weights and cross-correlation coefficients of a heterogeneous HORN^n^ as in A. Note the more decorrelated activity compared to a HORN^h^ already in the untrained state. **D**. Performance of different instances of a heterogeneous HORN^n^ (64 nodes) network as a function of training steps when trained with different learning rules. Learning rules as in B. Note the baseline (reservoir HORN) performance of the anti-Hebbian rule in this initially more desynchronized network (compared to the case in B) and the performance increase compared to baseline resulting from a Hebbian rule.

To investigate whether BPTT could be substituted by correlation-based learning, we implemented simple unsupervised additive (conventional) Hebbian rules as well as anti-Hebbian rules for the activity-dependent modifications of the recurrent connections (see Methods). As a performance baseline, we trained a HORN in which the recurrent connection weights were frozen, and only the input and readout connections were plastic (“reservoir HORN”). Subsequently, we allowed the recurrent connections to undergo Hebbian modifications. In the case of homogeneous networks, the application of a fully unsupervised anti-Hebbian rule resulted in a task performance close to that attained when the recurrent connections were trained by the fully supervised BPTT rule (Fig. 4B). The reason is that the anti-Hebbian rule first promoted selective desynchronization of the homogeneous network, allowing for the later build-up of stimulus-specific correlation structures. For heterogeneous networks that already express more decorrelated activity in the untrained state (Fig. 4C), the conventional Hebbian rule but not the anti-Hebbian rule increased performance above baseline (Fig. 4D). The reason is that the Hebbian modifications enhanced the already existing stimulus-specific correlation structure, whereas the anti-Hebbian rule could not induce further modifications of the already decorrelated activity patterns. In non-oscillating networks, BPTT training also resulted in weight distributions compatible with Hebbian principles in some cases, but to a lesser degree than observed for HORNs. We attribute the accelerated acquisition by HORNs of stimulus-specific weight distributions both in non-supervised and supervised learning to the unique dynamics of coupled oscillators. Their ability to engage in resonance amplifies correlated, synchronous activity patterns and makes them more sustained. These are ideal conditions for an unsupervised Hebbian mechanism that reinforces weight distributions and supports the evolution of stimulus-specific activity landscapes. Likewise, the BPTT algorithm also profits from the amplification through resonance of “meaningful” response constellations that support the orthogonalization of representations in the high-dimensional state space.

Taken together, these results provide a proof of principle that unsupervised Hebbian learning at the level of recurrent connections in HORNs supports the segregation of stimulus-specific dynamic states and thus facilitates their classification (Fig. 4B,D).

### Geometric input drives self-organization and traveling waves

To test how HORNs process simultaneous geometric input rather than time series, we trained networks that receive spatially organized input (Fig. 5A). To this end, we activated each node for one time step at an intensity corresponding to the sum of the intensity values of the MNIST pixels within its RF (Fig. 5A). We then gave the network 150 time steps to process the stimulus before performing a linear read-out (Fig. 5B). For training, we used the BPTT algorithm as before, started with an all-to-all random connectivity, and kept the weights of the input layer fixed, while the recurrent and readout weights were plastic. After training for 10 epochs, the classification accuracy on the test set was found to be around 90% (data not shown). When stimulated, each of the simultaneously activated nodes responded with a damped oscillation (Fig. 5B). These activities spread throughout the network and led to traveling waves and complex interference patterns. The direction and shape of the waves differed for responses evoked by different stimuli (Movies S1, S2). HORNs are able to sustain oscillatory activity in the form of a standing wave in each DHO node. These waves, once initiated, are sustained and give rise to global interference patterns. In contrast, traveling waves in RNNs without oscillating nodes interfere only when wave fronts collide.

**Figure 5:**
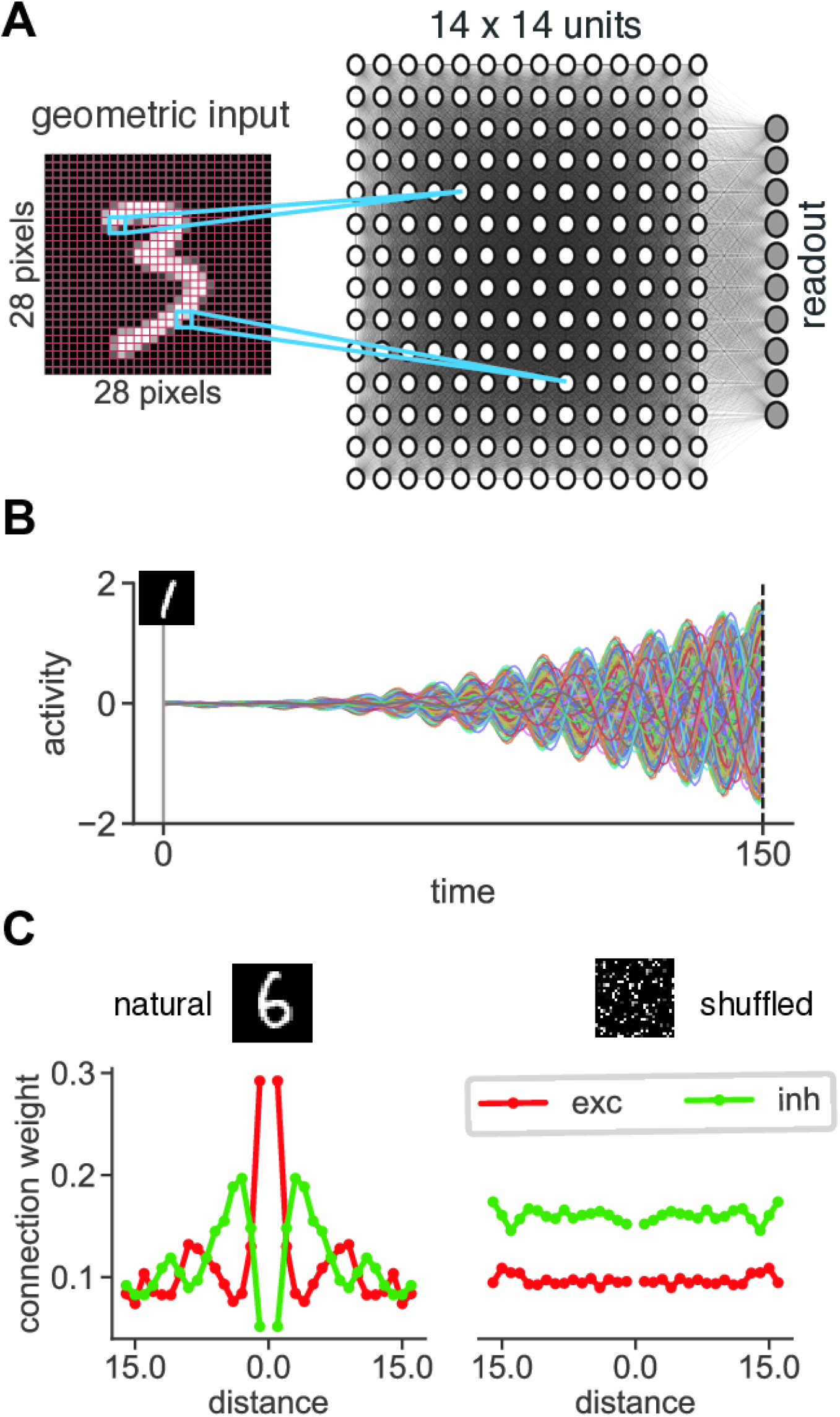
Self-organization in HORNs driven by geometric input. **A**. Schematic of geometric MNIST input provided to an all-to-all connected homogeneous HORN (196 nodes). Each DHO node receives input from its 2×2 pixel receptive field. Input weights are constant and fixed, recurrent and readout weights are trainable. Readout nodes are connected to the entire HORN. **B**. Example network dynamics of a HORN^h^ (196 nodes) stimulated as in A after training with BPTT for 2000 steps (achieved classification accuracy around 90% for both natural and shuffled stimuli). Input is flashed to the network during one time step at *t* = 1. Readout is performed after 150 time steps of network activity during which no input is given. Curves represent the activity of all nodes in the network (node identity is color-coded). Note the increasing amplitudes of reverbrating activity and the drifts in phase of the oscillatory responses. **C**. Distance-dependent connection weights of excitatory (red) and inhibitory (green) connections in HORN^h^ networks after training with BPTT for 2000 steps on geometric stimuli. Stimuli were either natural (left) or shuffled (right) MNIST digits. Note the Mexican-hat-like connectivity structure among nodes in networks trained on natural stimuli and the lack of structure in the connectivity in networks trained with shuffled stimuli.

In our model, nodes coactivated by a flashed stimulus exhibit stimulus-locked synchronized oscillations. As the BPTT algorithm mimics Hebbian plasticity in HORNs, this predicts that during learning these synchronously active nodes should increase their mutual coupling. Consequently, we observed a shift from unspecific all-to-all connections to spatially restricted connections with a distant-dependent decay of coupling strength, following a Mexican hat-like shape (Fig. 5C, left). This connectivity captures the essential structural feature of the MNIST stimuli, the continuity of their contours, and the spatial vicinity of activated nodes. During testing (recall) of the trained model, a particular stimulus again induces synchronized oscillations in a respective constellation of nodes, and, due to the enhanced coupling of those nodes in the trained network, these now engage in resonance which leads to an increase in response amplitude. As expected, such distant dependent connectivity patterns did not emerge when training HORNs on shuffled MNIST stimuli (Fig. S10), although the networks achieved a comparable classification accuracy on the test set (Fig. 5C, right). Here, other priors than spatial continuity and vicinity were installed in the architecture of the coupling connections, namely those representing the statistical contingencies of the shuffled MNIST digits without local spatial structure.

### Spontaneous and evoked activity

Cortical networks are spontaneously active and in sensory cortices, stimulation typically causes a reduction in variability (Fano factor) and leads to the emergence of stimulus-specific substates [73, 74, 75]. To model spontaneous activity in HORNs, we subjected each DHO node to random discrete jumps in its velocity by a given amount according to a Poisson process (see Methods). After training a spontaneously active heterogeneous HORN on sMINST, we analyzed network activity before, during, and after stimulation with sMNIST stimuli (Fig. 6). During stimulation, spontaneous activity was replaced by structured time series corresponding to the sMNIST digits. In this way, we mimicked protocols used in physiological experiments. Here, the unstructured resting activity of sensory organs, the main drive for spontaneous cortical activity, is temporarily replaced by stimulus-induced structured activity. Analysis of the activity of both single nodes and the entire network showed that the variance of activity was high before stimulation, decreased during stimulation, and then recovered to pre-stimulus levels after stimulus offset (Fig. 6A). Principal component analysis (PCA) revealed that the state space of spontaneous activity spans a large but confined space that comprised the subspaces of stimulation-induced stimulus-specific response vectors (Fig. 6B). The reason for this confinement of the spontaneous state is that the connectivity of the trained HORN is not random, but highly structured through learning (see “Learning Priors” above). This structure of the coupling connections, in turn, shapes the dynamics of the network. The temporal evolution of network dynamics in PCA space shows that network dynamics rapidly converges to a stimulus-specific substate once a stimulus is presented and that this specificity fades after stimulus offset (Fig. 6C). At the same time, this collapse onto such a substate is associated with a reduction in the variance of activity (Fig. 6A). Again, similar stimulation-dependent changes in network dynamics have been reported for natural networks [2, 76, 73].

**Figure 6:**
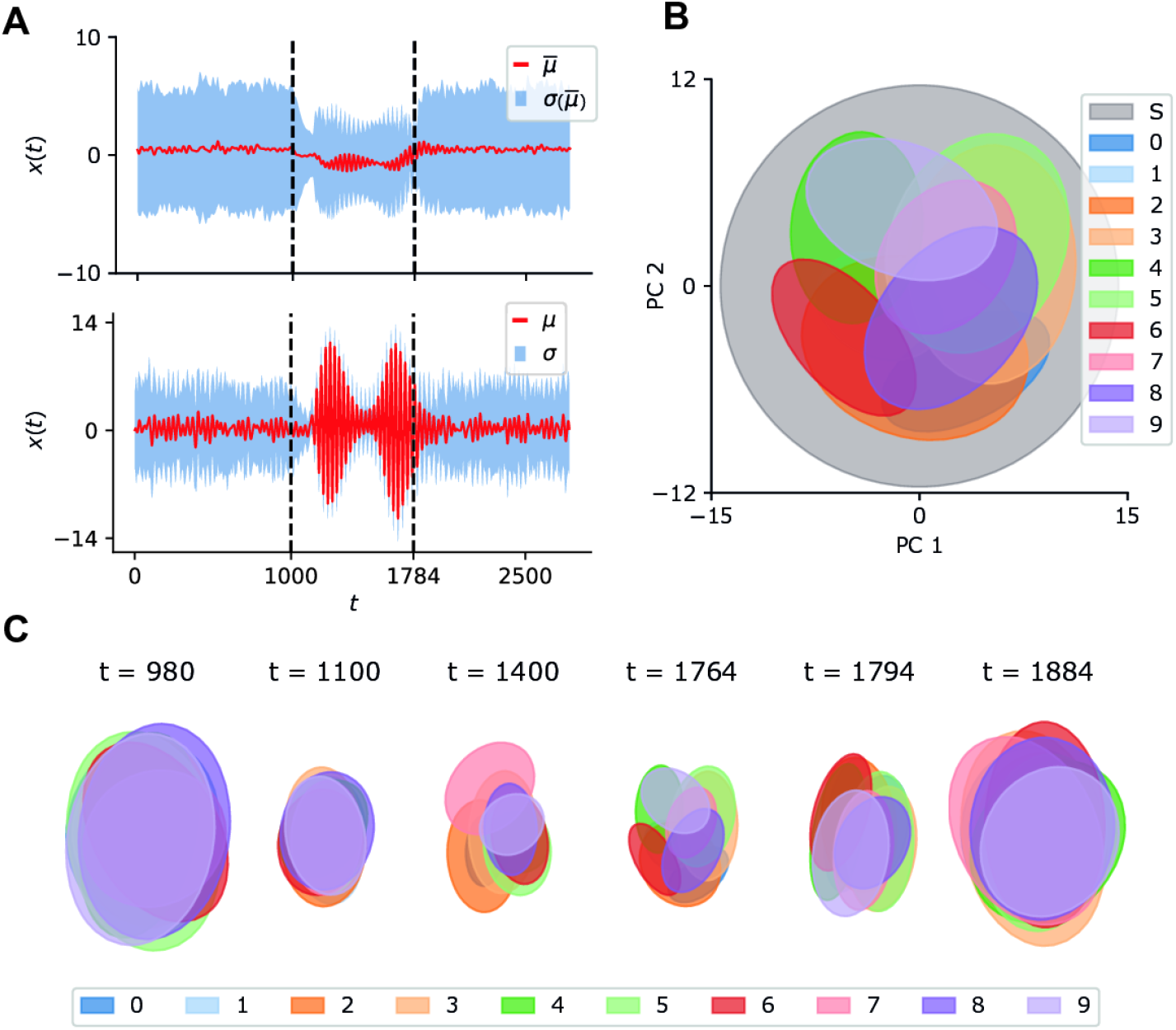
Spontaneous and evoked activity in a heterogeneous HORN network of 16 nodes trained on sMNIST. **A**. Mean and standard deviation of amplitude dynamics computed over 500 samples of the sMNIST digit 0. Top: Mean amplitude *ū* and standard deviation of mean amplitude *σ*(*ū*) calculated for the entire network. Bottom: Mean and standard deviation of amplitude dynamics of one selected DHO node. Dashed lines mark stimulus onset (*t* = 1000) and offset (*t* = 1784), respectively. **B**. Sliding-window analysis of PCs of network activity (window size: 20 time steps). Ellipses show 2*σ* confidence intervals, stimulus conditions are color-coded (S = spontaneous activity). **C**. Evolution of network dynamics in PC space under different stimulus conditions (*t <* 1000: spontaneous activity; 1000 ≤ *t* ≤ 1784: sMNIST stimulus presentation; *t >* 1784: spontaneous activity). Ellipses show 2*σ* confidence as in C.

## Discussion

### Controlled oscillations

The implementation of characteristic features of the mammalian cerebral cortex in simulated RNNs revealed a computational principle based on oscillatory activity. The most important step was to endow the network nodes with the propensity to oscillate. RNNs without oscillatory nodes can also produce oscillatory responses. However, such oscillatory activity is an emergent property of complex network interactions, often of transient nature, and hard to control. This makes it difficult to exploit these emergent oscillations for gradient-based learning schemes and assess their functional significance. For this reason, we opted for a synthetic approach and enforced oscillatory activity at each network node. This strategy allowed us to study the functional relevance of oscillatory dynamics and to identify the computational principle responsible for the increased performance of HORNs. This strategy also allowed us to establish close relationships with the dynamics of natural networks such as the cerebral cortex and assign a functional role to the oscillatory activity observed in this structure. Inspired by experimental findings [27, 77], HORN network nodes were configured as damped harmonic oscillators (DHOs) rather than leaky integrators or gated units, as used in other RNN architectures. This modification endowed both the single network nodes and the resulting networks with specific and easily controllable oscillatory dynamics. Individual nodes acquire the ability to extract features through resonance and, more generally, to modulate gain in a frequency-dependent manner. In turn, networks generate holistic transient stimulus representations characterized by wave interference patterns [78, 38, 39].

Testing performance on standard pattern recognition benchmarks revealed that a gradient-based learning scheme can capitalize on the extended dynamical repertoire of HORNs, leading to substantially enhanced performance relative to RNNs with non-oscillatory nodes. The improvements were manifest in learning speed, noise tolerance, and parameter efficiency. These findings of improved task performance are in line with previous studies in the field of machine learning that investigated RNNs with oscillating nodes [31, 38, 39]. The enforcement of oscillatory activity in each network node acts as an inductive bias in RNNs [55] and this bias can increase the practical expressivity of machine learning models. A famous example is the inductive bias introduced by convolutional architectures in feedforward neuronal networks that lead to convolutional neuronal networks that revolutionized deep learning [52].

The virtue of controlled oscillations for computations in RNNs has recently been demonstrated also in machine learning applications of spiking networks [79, 80]. Performance was enhanced when the subthreshold membrane potential was allowed to oscillate. This hints at a universal computational principle that is based on coupled oscillators and permits wave-based representations. Note that this principle need not be limited to neuronal populations, but could also act at the level of a single neuron [81].

When HORNs were trained with geometrically organized stimuli, they developed local connections between nodes. Such locally connected oscillator RNNs [38, 39] can be interpreted as a discretization of a neural field model with a specific connection kernel that implements a damped wave equation [82, 83]. These field models, formalized with partial differential equations, can be understood as a continuous approximation of HORN networks with specific local connection kernels. Similar arguments also hold for studies of traveling waves in RNNs without oscillating nodes [84]. Local oscillations and global waves are two sides of the same coin, and the dynamics of HORNs and field models are equivalent under certain conditions. To elucidate the intricate relationship between HORNs and neural field models is left for a future study.

### Reasons for increased performance

The good performance of HORNs is due to several reasons, and these are closely related to the propensity of network nodes to engage in oscillations. First, the ability of single network nodes to extract features from temporally modulated sensory signals through resonance contributes to the noise resilience of HORNs. “Innate” preferences for stimulus features (controlled by the parameters *ω, γ, α*) allow individual nodes to efficiently extract and encode stimulus features already in an untrained network. If input signals lack temporal structure, the oscillatory properties of the nodes are still beneficial because they transform sustained inputs into oscillatory responses. This transformation allows computations in the common format of temporally modulated signals which prevail in the communication among nodes.

Second, the comparison of learning speed in oscillating and non-oscillating networks indicates that oscillations facilitate both BPTT and Hebbian learning. Both strategies installed priors that were stored in the weight distributions of the recurrent connections and shaped the dynamics of stimulus representations in a way that facilitated discrimination by an affine readout. The likely reason for faster learning of oscillatory networks is that nodes responding to related stimulus features tend to synchronize and engage in resonance, in particular, once learning has started or if priors have been installed previously in the recurrent connections. Although the correlation-sensitive Hebbian mechanism was expected to profit from prolonged temporal correlations and synchrony, it is less obvious how the BPTT algorithm exploits oscillatory activity (see [55] for a detailed analysis). The discretization scheme used for the oscillator differential equations introduces temporal residual connections that stabilize gradient propagation in BPTT learning (see Methods and [31, 55]). Furthermore, the oscillating dynamics of each DHO node temporally modulate gradients, changing the practical expressivity of the networks (see Methods, Fig. S11 and [55]).

Third, DHO units in a HORN network collectively process stimuli in a fully distributed manner by converting sensory input into waves. Initially, these are standing waves in each oscillator, but then they spread and give rise to complex interference patterns on the network level [16, 17], findings that are compatible with physiological evidence [78]. This representation provides a coding space of massive dimensionality, and most importantly, it permits the superposition of information about multiple spatially and temporally segregated events. This allows HORNs to analyze and encode simultaneously not only spatial but also temporal relations between a large number of stimulus features and to generate holistic representations of the correlation structure of complex input constellations.

Note that RNNs without oscillating nodes also generate traveling waves [85] and that it is conceivable that these networks can also learn to exploit these waves for computations. However, the computationally relevant interference required for the coding of relations between different stimuli is restricted in space and time to the collision of wavefronts. Testing the functional role of traveling waves will likely require the same control strategy that we applied in this study to determine the function of oscillations.

### The virtues of heterogeneity

Heterogeneity improves the performance of RNNs because it increases the dimensionality of the state spaces of the networks. Having oscillatory nodes allowed us to increase heterogeneity also in the temporal domain by varying preferred oscillation frequencies. As expected, this increased task performance. Another advantage of heterogeneity in the temporal domain is that it brings network dynamics closer to criticality [86, 60, 55, 61]. Dynamics close to criticality are a hallmark of cortical networks and provide computational benefits summarized in the “critical brain” hypothesis [87]. These benefits are due to the emergence of long-lived transient and metastable states [62, 88]. HORNs also encode information in transients. Therefore, they are capable of coding with sequences of metastable states and ghost attractors [89, 90] if in a regime close to criticality. This distinguishes their dynamics from that of attractor networks [91, 92]. In the latter, critical slowing down limits computational power near criticality [93]. More studies are needed to better understand these transient dynamics in HORNs and to identify related activity in biological networks [89, 90].

In addition to enhancing heterogeneity by varying the preferred oscillation frequencies of the nodes, we induced heterogeneous conduction delays to deliberately induce phase shifts. This further increases the dimensionality of the networks’ coding space, which can be exploited for computation.

The implementation of these physiologically plausible features typically increased performance, while not increasing the number of trainable system parameters. It endowed even untrained networks with sensitivity to a broad range of correlation structures that could then be exploited by subsequent learning to accelerate learning speed. Most importantly, heterogeneity improved the ability to process stimuli with novel characteristics. This alleviates the need for fine-tuning and the search for optimal stimulus-specific parameters. Such fine-tuning processes are cumbersome and computationally expensive in artificial systems and are likely not realizable in neuronal systems. The gain of function by heterogeneity was particularly pronounced in larger networks, suggesting that one-shot learning, a hallmark of biological systems, is facilitated in very large, heterogeneous networks.

The advantage of nodes with a propensity to oscillate and to introduce heterogeneity by fine-tuning the preferred oscillation frequency is documented also by the simulations of two-layer networks. These networks showed enhanced performance at the same number of parameters, in particular when the higher layer operated at lower preferred frequencies than the lower layer. This allows the multi-layer network to operate in different frequency bands and to perform parallel analyses of input patterns at different temporal scales in each layer. This finding was the result of a grid search for optimal parameter settings and shares similarities with the organization of the cerebral cortex. Here, too, oscillation frequencies decrease as one progresses from lower to higher processing levels [10, 94]. Slower oscillations at higher levels can establish relations among temporally segregated stimuli over longer time intervals, which could support chunking.

Interestingly, according to basic physics, waves with slower frequencies tend to travel over longer spatial distances, in our case over a larger number of network nodes. In the cerebral cortex, higher areas integrate information from increasingly diverse and spatially remote processing streams, as reflected by their large, often polymodal, and multiselective receptive fields. Assuming a wave-based presentation [78], operating at lower oscillation frequencies would allow these higher areas to integrate information over larger temporal and spatial scales, favoring holistic processing of information and multimodal binding.

Taken together, we propose that the heterogeneity of preferred oscillation frequencies and coupling delays in natural neuronal systems is likely not a reflection of nature’s imprecision but an efficient solution to computational challenges. Heterogenity increases the dimension of the available coding space, enables the robust processing of noisy sensory stimuli with highly varying spectral characteristics, alleviates the computationally challenging problem of finding optimal system parameters, and allows networks to exploit the computational advantages of operating close to criticality.

### Relations to neurobiological systems

HORNs reproduced several characteristic features of the dynamics and organization of natural neuronal systems, in particular of the cerebral cortex and probably also the hippocampus. In addition, the simulations allowed us to assign concrete functions to features of natural networks whose role in information processing is still a matter of discussion.

Our simulations show that learning-dependent complex, transient, and stimulus-specific synchronization patterns provide benefits for information processing and identify as an underlying mechanism the oscillatory properties of network nodes. This supports the hypothesis that oscillations and synchrony, which are also observable in neuronal systems [95] are functionally relevant and not epiphenomena.

In most of our simulations, we used time series as input signals. One might argue, therefore, that our simulations capture only the temporal relations in sensory signals and neglect spatial relations that also play a prominent role in the processing of sensory information, especially in the visual system.

However, simulations with geometrically organized input patterns yielded results very similar to those obtained with time series data. Thus, the identified computational principles can handle spatial and temporal relations among input signals similarly and represent computation results in the same format. This benefits computations in sensory cortices receiving both temporally and spatially structured input, aiding cross-modal and inter-areal communication. For both temporally and spatially structured stimuli, learning led to synaptic weight configurations that decay with distance and capture the Gestalt criteria of continuity and vicinity, a property also known from neuronal systems [2, 76]. In the visual cortex, the basic layout of recurrent connections is genetically determined, but experience-dependent pruning of these connections further enhances their selectivity through a Hebbian mechanism [96].

Stimulation of locally connected HORNs led to traveling waves that resemble closely those observed in natural neuronal networks [16, 78]. Traveling waves are also a hallmark of oscillatory RNNs in which local connectivity was enforced by design [38, 39]. Wave-based representations allow for very high-dimensional representations and manifold coding strategies. Consequently, numerous hypotheses have been proposed regarding the functional role of traveling waves [17, 81]. In a wave-based model of motor cortex [81] the direction and wavelength of traveling waves are used to structure commands in a way that is easily decodable by the dendritic arbors of neurons in the descending motor system. However, the exact function of traveling waves in the sensory cortices is still not fully understood.

Another similarity between the dynamics of HORNs and the cerebral cortex is the temporal evolution of responses in simulations with geometrically structured stimuli. The initial transient responses were amplified by reverberation, increasing the decodability of the dynamic state due to better segregation of stimulus-specific principal components of the population vector [76]. This state can be seen as a highly parallelized search for the best match between sensory evidence and learned priors [2]. Thus, one of the core functions of predictive coding, the matching of sensory evidence with stored priors, can be realized through self-organizing dynamic interactions in oscillatory recurrent networks.

During learning, nodes activated by semantically related features increase their mutual coupling, and during recall these nodes self-organize into a stimulus-specific assembly with synchronized and jointly enhanced responses. This dynamic association of nodes is also observed in the visual cortex for neurons tuned to perceptually bound features [11, 76] and is at the core of the binding by synchrony hypothesis (BBS) [20]. HORNs, by exploiting the resonance properties of coupled oscillators, reproduce this important feature of natural cortical networks.

Close similarities with the dynamics of natural cortical networks were also apparent in spontaneously active HORNs. Stimulation reduced the variance of spontaneous activity [73] and induced a transient convergence of the network dynamics to stimulus-specific substates, as predicted in [2]. These substates occupy a region within the subspace formed by spontaneous activity and are the result of a comparison between sensory evidence and priors stored in the weight distribution of feed-forward and recurrent connections. Thus, spontaneous activity can be considered as a superposition of fragments of learned stimulus-specific representations.

In addition to the aforementioned reproduction of many physiological phenomena, additional physiological experiments can now be designed to examine specific predictions derived from the present study. These experiments will require massive parallel recordings of neuronal activity both within and across cortical areas with high spatial and temporal resolution to capture the spatio-temporal dynamics of traveling waves and their resulting interference patterns.

### Concluding remarks

Taken together, the present results not only unveil the computational principles accessible to HORNs and other oscillator networks, but also allow for a functional interpretation of numerous experimentally verified physiological phenomena whose roles in information processing so far have been elusive or have caused controversial discussions. Plausible functional roles can now be assigned to (i) the propensity of nodes to oscillate and the resulting dynamical phenomena such as synchronization, desynchronization, resonance, entrainment, and traveling waves [12, 13, 14, 15, 17, 18]; (ii) the diversity of preferred oscillation frequencies, their non-stationarity and context dependence [3, 4], (iii) the heterogeneity of the conduction velocities of the recurrent connections [5, 6], (iv) the decrease of oscillation frequencies in higher areas of the cortical processing hierarchy [10, 94], (v) the Hebbian adaptivity of recurrent connections [7, 8]; (vi) the emergence of context-dependent dynamic receptive fields by network interactions [68, 70], and (vii) the reduction of variance in network dynamics during stimulus presentation [73]. The simulations also suggest a physiologically plausible scenario for the rapid and parallel matching of sensory evidence with stored priors through self-organized convergence of network dynamics to classifiable, stimulus-specific, dynamic substates. These substates consist of highly structured, high-dimensional dynamical landscapes that unfold due to interference of wave patterns in amplitude, frequency, and phase space. In essence, the described networks of coupled oscillators perform highly parallelized analog computations in high-dimensional state spaces. This allows networks to simultaneously relate a large number of spatially and temporally structured input variables, a capacity ideally suited to accomplish context-dependent feature binding and scene segmentation. Accordingly, attempts are made to exploit the principle described in this study in machine learning architectures designed to perform scene segmentation [97]. Moreover, the computational strategy implemented by HORNs is also well suited to overcome challenges that require the simultaneous evaluation of multiple nested relations. Typically, such problems need to be solved in language comprehension. Interestingly, biological systems are at ease with solving the binding problem, with the segmentation of cluttered scenes and with the analysis of complex time series (e.g. spoken language) while these tasks are notoriously difficult for digital computer architectures that typically rely on serial feedforward processing. Therefore we believe that nature solves such hard problems by means of analog computations of the kind described in this study. We predict that it will be possible to implement the computational principle presented here in analog hardware that runs at room temperature, is miniaturizable, and highly energy efficient. In combination with novel electrical elements mimicking Hebbian synapses such as memristors, this novel principle will likely enable the design of self-adapting devices for machine learning applications that can ideally complement existing digital technologies.

## 1 Materials and Methods

### DHO nodes

We model each HORN node as a driven damped harmonic oscillator (DHO) with feedback, defined by the second-order ordinary differential equation (ODE)

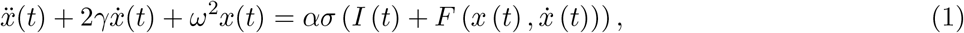

with *x*(*t*) denoting the oscillator amplitude, *γ >* 0 the damping factor (Fig. S1A), *ω >* 0 the natural angular frequency (Fig. S1B), and *α >* 0 an excitability parameter. *σ*(·) denotes an input nonlinearity that models local resource constraints and is chosen as tanh(·). *I*(*t*) denotes a time-varying external input and 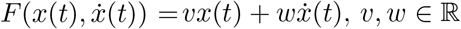 the feedback input (see below).

Depending on the quotient *s* = *ω/γ*, the solutions of (1) have qualitatively different relaxation dynamics (Fig. S1A,B): (i) When *s >* 1, solutions are given by oscillations with decaying amplitude (subcritical damping). (ii) When *s* = 1 or *s <* 1, solutions are given by non-oscillating exponentially decaying curves (critical damping and supercritical damping, respectively).

To define a discrete-time update equation for a DHO node, we first rewrite (1) as a system of two first order ODEs

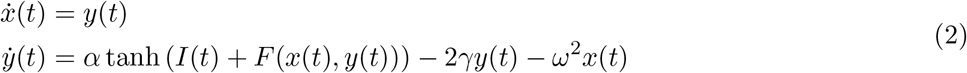

and then discretize (2) using a microscopic time constant *h* to obtain the update equation for a discrete-time HORN network in which each node performs an Euler integration of (2) at each time step. As an explicit Euler integration scheme can be numerically unstable when used for the integration of (2), we implemented a symplectic Euler integration scheme [98] for each node and obtain the update equations

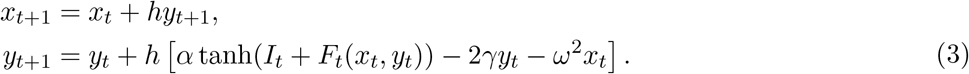

Here, *t* denotes a discrete discrete time index 1 ≤ *t* ≤ *T, I*_*t*_ is the external input at time *t*, and *F*_*t*_ the feedback input of the node to itself. The initial conditions are chosen as *x*_1_ = *y*_1_ = 0 unless otherwise noted. The natural frequency *ω* corresponds to a period *ω*_*p*_ = 2*π/*(*ωh*) and the memory half-life time is proportional to 1*/*(*γh*), both measured in input time steps.

The microscopic time constant *h* defines a scaling factor of the input time to the system time. It can be interpreted as the sampling frequency of the system, and the linear Nyquist frequency is *h*_Nyquist_ = 1*/*2*h*. Therefore, the theoretical upper limit of the natural (circular) frequency parameter *ω* for any DHO node must be *ω*_max_ *< π/h*. In practice, this bound is lower as the Euler integration step in (3) becomes numerically unstable (data not shown).

Since the number of integration time steps always equals the length *T* of the input time series, the numerical integration of the system (3) is invariant under changes in *h* → *h*^′^ when system parameters are rescaled according to

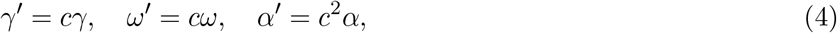

where *c* = *h/h*^′^. To allow a better interpretability of the natural frequency parameter, we set *h* = 1 without loss of generality to link the two time scales and express the natural frequency *ω* in period nodes of the input as *ω*_*p*_ = 2*π/ω* when convenient. A value of *h* = 1 will be assumed throughout the manuscript if not stated otherwise.

### HORN networks

A HORN network is a system of *n* additively coupled DHOs of the form (3). Given a time-varying external input *S*_ext_ = (**s**_1_, …, **s**_*T*_), **s**_*t*_ ∈ ℝ^*m*^, the update equations of a HORN are

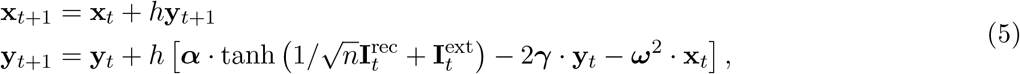

where 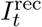 and 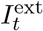 denote the recurrent and external input to each node at time t, respectively, defined as

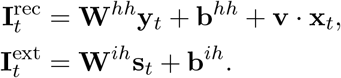

Above, lowercase boldface symbols denote vectors of length *n*, uppercase boldface symbols denote matrices, · denotes element-wise multiplication, **W**^*hh*^ is the *n* × *n* matrix of recurrent coupling weights, **W**^*ih*^ is the *n* × *m* matrix of input weights, and **b**^*hh*^, **b**^*ih*^ are bias vectors. As for the single-node case, the update equations implement a symplectic Euler integration scheme. The feedback input from a node to itself *F* (*x*_*t*_, *y*_*t*_) is *F* (*x*_*t*_, *y*_*t*_) = *vx*_*t*_ + *wy*_*t*_ with 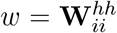. Note that the values of the parameters *ω, γ, α* can vary between DHO nodes. To train a HORN on a task, an affine readout of the network state 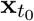 at time *t*_0_ is defined as a map 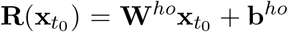, where **W**^*ho*^ is the *k* × *n* readout matrix and **b**^*ho*^ is a bias vector. As previously, we will set *h* = 1 to link the input and system time scales, unless otherwise stated. See below for a description of network simulations.

### Other RNN architectures

A simple Elman RNN [99] with a linear readout (referenced by the label “tanh” throughout all plots) was implemented by the update equation

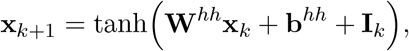

where the input **I**_*k*_ is defined as above. Similarly, a “leak” RNN consisting of leaky integrators was implemented by the update equation

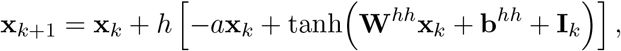

where *a* is a leak constant that controls memory decay, see also [100] in which *a* = 1, *h* = 1. For the LSTM and GRU based RNNs, the implementations of LSTM and GRU nodes provided by PyTorch (version 1.9.0) [101] were used; see below.

### Number of trainable parameters

Performance comparisons for different RNN architectures were carried out at the same number of trainable parameters. Assuming an RNN with *n*_*h*_ nodes receiving input of dimensionality *n*_*i*_ through an affine projection layer, and a readout to an *n*_*o*_-dimensional output vector via an affine readout layer, the number of trainable parameters is *N* = *c*(*n*_*i*_ + *b*_*i*_)*n*_*h*_ + *cn*_*h*_(*n*_*h*_ + *b*_*h*_) + (*n*_*h*_ + *b*_*o*_)*n*_*o*_, where *c* is the number of state variables per node (*c* = 3 for GRU, *c* = 4 for LSTM, *c* = 1 for the simple and leak RNNs) and *b*_*i*_, *b*_*h*_, *b*_*o*_ is either 1 (bias term present) or 0 (no bias term present). HORNs also have one state variable per node (i.e. *c* = 1), but an additional amplitude self-connection term for each node, leading to *N*_HORN_ = (*n*_*i*_ + *b*_*i*_)*n*_*h*_ + *n*_*h*_(*n*_*h*_ + 1 + *b*_*h*_) + (*n*_*h*_ + *b*_*o*_)*n*_*o*_. To construct a network of a given architecture with a given number *N* ^*^ of trainable parameters, we solve the quadratic equation *N* = *N* ^*^ after plugging in the known values for *n*_*i*_, *n*_*o*_, *b*_*i*_, *b*_*h*_, *b*_*o*_ and taking the integer part of the positive solution to obtain the largest number *n*^*^ of nodes so that the number of trainable parameters does not exceed *N* ^*^. See Fig. S19 for the scaling behavior of trainable parameters as a function of the number of nodes for different RNN architectures for *n*_*i*_ = 1, *n*_*o*_ = 10, *b*_*i*_ = *b*_*h*_ = *b*_*o*_ = 1.

### Datasets

The **MNIST** data set [102] of hand written digits consists of a training data set of 60,000 samples and a test set of 10,000 samples which was used to assess classification performance (measured by classification accuracy) of different networks. Each sample is given by a 28×28 matrix of intensity values between [0, 255] which are scaled to [0, 1]. Here, two variants of the MNIST data set are considered, the sequential MNIST (sMNIST), and the permuted sequential MNIST (psMNIST). For sMNIST, the 28×28 intensity values of a sample are collected in scanline order from top left to bottom right to form a time series of length 784 that is then fed to the networks step by step. For psMNIST, a random but fixed permutation on 784 elements is applied to the stimulus before feeding the input to the network. This removes the dominant low frequency structure present in sMNIST digits and distributes stimulus information across all frequencies (Fig. S10, S5). Noise was added to the sMNIST or psMNIST time series in form of i.i.d. additive white Gaussian noise at the pixel level and sampled from *N* (0, *σ*^2^). Stimulus values were clamped to [0, 1] after noise was added.

The **Line segments data set** (LSDS) consists of samples of 28×28 pixel intensity values with one or more line segments of a given angle. The data set has four parameters that define the nature of its samples: The number of angles *n*_*a*_, the maximal number of segments per sample *n*_*s*_, the minimal and maximal length of the line segments *l*_min_ and *l*_max_. Additionally, it comes in two flavors: with random segment locations or centered segment locations. To construct one sample, first each entry of a 28 × 28 pixel canvas is set to 0 and an angle 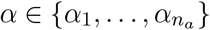 is chosen at random with *α*_*i*_ = (*i* − 1)*π/n*_*a*_, i.e. each sample only shows line segments of the same angle. Then the number of line segments *k* to be shown is chosen as a random integer in 1, …, *n*_*s*_. For each line segment *i*, first a random integer length *l*_*i*_ is drawn from *l*_min_, …, *l*_max_. Then a line segment of length *l*_*i*_ pixels is drawn on the 28 × 28 canvas with intensity 1.0 (using the Python package opencv-python, version 4.6). If the line segments are to be centered, they are drawn so that their central point coincides with the center of the 28 × 28 canvas. If the locations of the segments are random, a random starting coordinate from the set {0, …, 21} is chosen for both the horizontal and vertical starting point of the line segment. The label for each sample is the angle *α* present in the sample and the networks are trained on a *n*_*a*_-class classification problem of predicting the angle shown in a given sample. We will write LSDS(*n*_*a*_, *n*_*s*_, *l*_min_, *l*_max_, *L*) with *L* ∈ r, c to refer to the data set with random (*L* = r) and centered (*L* = c) segment locations. We use two versions of the LSDS in the study, LSDS(32,5,11,c) which we will refer to as LSDSa, and LSDS(10,3,8,24,r) which we will refer to as LSDSb. Samples of line segment data sets are shown in Fig. S16.

The **EMNIST** (balanced) data set [103] consists of 131,600 handwritten characters belonging to 47 classes, representing handwritten digits as well as uppercase and lowercase Latin characters. Each character is represented by a matrix of 28×28 pixel intensity values as for MNIST. The training data set is of size 112,800, the test set of size 18,800. EMNIST samples have the same format as MNIST, and their signal properties as time series are similar to those of MNIST digits.

The of **Spoken Digits Data Set** (SDDS), was chosen as a subset of the Speech Commands data set (version 0.0.2) [104] (SCDS). The SCDS consists of 65,000 audio audio recordings (1 second length, mono, 16kHz sampling rate) of spoken words and digits obtained from thousands of people and spans 30 different classes. The SDDS data set was formed by choosing the 10 classes from the SCDS representing spoken digits 0 to 9, forming a data set of 23,666 samples (with each class consisting of around 2730 samples). Of the SDDS, a random selection of 10% of all samples was used as the test set, and the rest was used as the training set.

To construct the input time series to networks, each spoken digit was first sub-sampled to 4000 Hz using the Python library librosa, followed by a normalization of the amplitude to the interval [−1, 1], performed per sample.

The power spectral densities of the time series calculated on samples from each data set is shown in Fig. S5.

### Simulations

All simulations were performed in PyTorch (version 1.9.0) [101]. All networks consisted of three trainable affine layers (input, recurrent, readout). For the Elman, leak, and HORN networks, custom implementations were used and the code is available at https://github.com/. For the LSTM and GRU networks, the standard implementations provided by PyTorch were used.

The initial weights and biases of all affine layers were randomly drawn from a uniform distribution 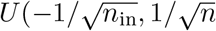 where *n*_in_ denotes the input dimension of the layer (default implementation in PyTorch torch.nn.Linear).

All parameters were learned with the BPTT algorithm. As loss function for the studied stimulus classification problems we used the cross-entropy loss function (torch.nn.CrossEntropyLoss). Weight updates were performed according to the AdamW optimizer with default settings (betas=(0.9,0.999), eps=1e-8, weight_decay=0.01) using a batch size of 128 for all data sets, unless otherwise stated. For the SDDS data set, we used a batch size of 64.

For each model, a hyperparameter grid search was performed using a network with 10^4^ trainable parameters to find the parameter values that resulted in a network with the highest classification accuracy on the test set after 10 training epochs. The parameters contained in the grid search were different per model and are described below. For all models, the grid search included the learning rate lr ∈ {10^−5^, 5 · 10^−5^, 10^−4^, 10^−3^, 5 · 10^−3^, 10^−2^}. Optimal learning rates for HORNs and the leak RNN were found to be 10^−2^ except for SDDS where they were 5 · 10^−3^. Optimal learning rates for the gated (LSTM, GRU) RNNs were determined to be 10^−3^ for all data sets (SDDS: 10^−4^). Learning rates for the Elman RNN were set to 10^−4^ for all data sets (SDDS: 10^−5^). Gradient clipping (*pytorch.nn.utils.clip_grad_norm*) was applied in the case of the Elman, GRU, and LSTM networks, as without gradient clipping these architectures showed unstable learning trajectories or failed to depart from chance-level classification performance for sMNIST. For these networks, the grid search included the learning rate and the clipping values *c* ∈ {0, 0.1, 1, 5, 10, 100}. A clipping value of 1 was chosen for all data sets. For the leak RNNs, the grid search included the learning rate, the time constant *h* ∈ {0.01, 0.05, 0.1, 0.2, 0.3, 0.4, 0.5} and the leak constant *a* ∈ {0.05, 0.1, 0.2, 0.3, 0.4, 0.5, 0.75, 1.0}. See Table S1 for the parameter values for each data set used. For homogeneous HORNs, the grid search included the learning rate, the time constant *h* ∈ {0.01, 0.05, 0.1, 0.2, 0.3, 0.4, 0.5}, the natural frequency *ω* ∈ {0.1, 0.5, 1.0, 2.0, 3.0, 4.0, 5.0, 10.0}, and the damping factor *γ* ∈ {0.001, 0.005, 0.01, 0.05, 0.1, 0.25, 0.5}, setting *α* = 1.0. For heterogeneous HORNs, as a first step the spread of values *h, ω, γ* among the best 10 performing homogeneous HORNs from the previous step were calculated as [*h*_min_, *h*_max_], [*ω*_min_, *ω*_max_], [*γ*_min_, *γ*_max_], respectively. Subsequently, each node of a HORN^*n*^ was randomly assigned parameter values *h*_*i*_, *ω*_*i*_, *γ*_*i*_ sampled in an i.i.d. manner from *U* ([*h*_min_, *h*_max_)), *U* ([*ω*_min_, *ω*_max_)), *U* ([*γ*_min_, *γ*_max_), respectively, where *U* ([*a, b*)) denotes the uniform distribution over the half-open interval [*a, b*).

See Table S.1 for dataset-dependent parameters for the HORN and leak networks. For HORN networks, the normalized parameters with *h* = 1 are given. For the implementation details of HORN networks with synaptic delays and multilayer networks, see below.

### Gradient stability

Given a loss function *L*, the gradients of *L* with respect to a recurrent weight *w* can be computed as [105]

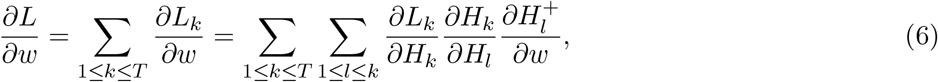

where *H*_*t*_ is hidden state of the network at the time *t, T* is depth of recurrent network (i.e. the number of input time steps), 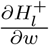 is the immediate derivative and where

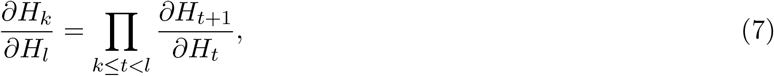

by iterated application of the chain rule. The propagation of the gradient at time *t* is defined by the be-havior of the Jacobian 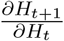. The vanishing gradient problem is characterized by 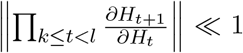, and 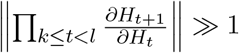 is called exploding gradient problem.

To study the dynamics of gradients in HORNN networks, we first make the following substitutions in the expression (5):

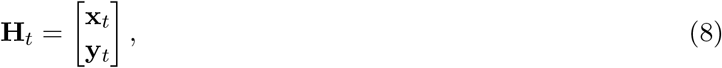

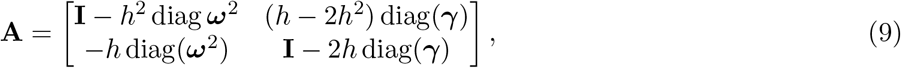

and denote by

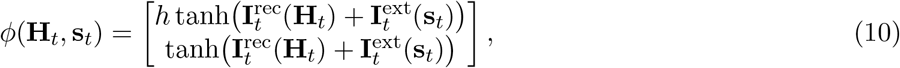

the transformation defined by the tanh nonlinearity, where **s**_*t*_, 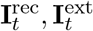 are as in (5), **I** denotes the identity matrix, and diag(**x**) the diagonal matrix with diagonal entries from the vector **x**.

These substitutions allow us to write the recurrent update step (5) as

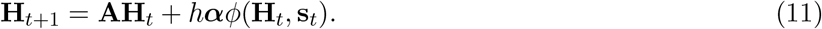

The sufficient condition to avoid both vanishing and exploding gradients is 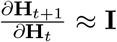. This condition can be achieved in (11) by choosing an Euler discretization time step *h* ≪ 1, or, equivalently choosing sufficiently small values for *ω, γ, α*. The scaling relationships for the system parameters *ω, γ, α* that result in equivalent system behavior under changes of *h* is described in Equation (4). Similar arguments hold for the RNN composed of leaky integrators, explaining their increased performance compared to a simple Elman RNN. In essence, the Euler integration scheme introduces residual connections between subsequent steps of HORNN dynamics, which ensure stable gradient propagation [106].

### Oscillating gradients

*L*^∞^ gradient norms of 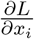 were computed based on network activity of a homogeneous HORN (64 nodes) obtained on the test set of sMNIST [107]. Gradients were computed for each time step of network activity (Fig. S11, right). Such modulated gradient magnitudes implement an attention mechanism that allows networks to pick up on certain spectral properties of stimuli (such as prominent frequencies) while ignoring others.

### Phase locking value

To analyze the pairwise synchronization between DHO nodes, we used the analytical representation of the amplitude signal *x*(*t*) which can be computed by means of the Hilbert transform as *X*(*t*) = *x*(*t*) + *iH* (*x*(*t*)), where *H*(·) is the Hilbert transform and *i* is the imaginary node. Then, the instantaneous phase *ϕ*(*t*) and the instantaneous magnitude *A*(*t*) can be computed as *S*(*t*) = *A*(*t*) exp (*iϕ*(*t*)). As a measure of nonlinear interactions, we use the pairwise phase locking value (PLV) calculated over the time window *T*, where *T* denotes the length of the input signal [108]. The PLV for two signals *x*_1_ and *x*_2_ in analytical representation is defined as

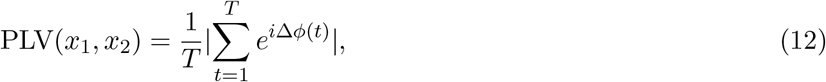

where Δ*ϕ*(*t*) = *ϕ*_1_(*t*) − *ϕ*_2_(*t*).

The phase locking value PLV takes values on the real interval [0, 1]. The case of PLV(*x*_1_, *x*_2_) = 1 indicates fully phase-locked synchronization of *x*_1_ and *x*_2_, while a value of PLV(*x*_1_, *x*_2_) = 0 indicates fully asynchronous signals *x*_1_, *x*_2_.

### PCA dimensionality of network activity

To measure PCA dimensionality, network activity vectors were collected using 1000 samples from the sMNIST test dataset. The activity vectors for all 784 time steps of stimulus presentation were treated as independent samples, resulting in 1000 · 784 samples. PCA was performed on the resulting dataset (Python package scikit-learn, sklearn.decomposition.PCA). As a result, every principal component constitutes the activation vector, which explains the amount of variance defined by the corresponding eigenvalue of the covariance matrix. The number of PCA components explaining 99% variance of the network’s activity was taken as measure of PCA dimensionality. PCA dimensionality measures the linear dimensionality and takes integer values on the interval [1, *N*], where *N* is the number of nodes in the network. To show the dynamics in PCA dimensionality over learning, it was computed for a sequence of intermediate instances over training for each network.

### Decoding of HORN activity

To decode the stimulus based on RNN activity, activity vectors were collected at each time step *t* of the stimulus presentation phase (*t* = 1, …, 784 for sMNIST), using 10000 samples from the test dataset. For each time step *t*, the resulting set of activity vectors was randomly split into a training set and a test set using a 80%/20% ratio. Using this dataset, an SVM classifier was trained on the 10-class stimulus classification problem for each time point *t* (Python package scikit-learn, sklearn.svm.LinearSVC). A regularization constant of *C* = 0.1 was chosen to ensure the convergence of all SVM classifiers. The confusion matrices were calculated using sklearn.metrics.confusion_matrix. Distance to the SVM decision hyperplane was calculated with sklearn.svm.LinearSVC.decision_function.

### Receptive fields

For each DHO node, the intrinsic receptive field (IRF) was calculated as the gain curve of the node [109]. The gain curve is computed using harmonic (sinusoidal) inputs. It is defined as the quotient between the DHO node amplitude, i.e. the stationary forced oscillation, and the input amplitude as a function of the input frequency *ω*_*i*_. For the intrinsic receptive field, these gain curves were calculated for an input amplitude of *A* = 1 over the time of *T* = 50000 time steps to allow the transient dynamics to settle.

The stimulus-dependent effective receptive field (ERF) of each DHO node in a HORN network was calculated as the mean most strongly driving stimulus (as the mean absolute amplitude computed over the entire presentation time). For this, network activity was first recorded for *n* = 10000 samples in a test set, and then the average of the 500 stimuli that resulted in the highest activation of a given DHO node was calculated to form the effective receptive field of a DHO node.

### Networks with synaptic delays

To implement synaptic conduction delays in a network of *n* nodes, we first defined a maximum delay *d*_max_ present in a network (measured in the number of discrete time steps) and subsequently assigned to each connection from a node *j* to a node *i* a delay of *d*_*ij*_ time steps. We then modified the recurrent input 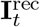 of the HORN update equation (5) to

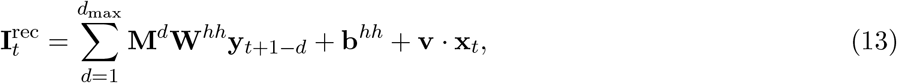

where **M**^*d*^ ∈ {0, 1}^*n*×*n*^, 1 ≤ *d* ≤ *d*_max_ are binary masks and the value of 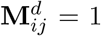 iff the connection from node *j* to node *i* has a delay of *d*, and 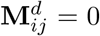 otherwise. Furthermore, we assume **y**_*i*_ = 0 for all *i <* 0. This approach allows us to continue to use the dense gradient methods implemented in PyTorch while effectively performing operations on *d*_max_ sparse matrices in each time step.

### Multi-layer networks

To implement a two-layer network with per-layer connectivity, we first choose the number of nodes per layer as *n*_1_ and *n*_2_. Furthermore, we introduce connection probability parameters *p*_1_, *p*_2_, *p*_F_, *p*_B_ where *p*_1_ and *p*_2_ model the recurrent connection probabilities within the layer 1 and 2, respectively, *p*_F_ is the feedforward connection probability from layer 1 to layer 2, and *p*_B_ is the feedback connection probability from layer 2 to layer 1. We set *p*_1_ = *p*_2_ = 1 unless otherwise noted, i.e. we choose fully connected layers. We then model the two-layer network by simulating a HORN network of *n* = *n*_1_ + *n*_2_ nodes, where the first *n*_1_ nodes represent layer 1, and the rest of the nodes layer 2. To model layer-specific connectivity patterns, we modify the HORN update equation (5) using masks as above and write

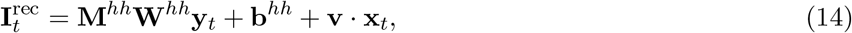

where the masking matrix **M**^*hh*^ ∈ {0, 1}^*n*×*n*^ has a block structure and defines the connections present in the network. The entries of **M** are sampled from a binomial distributions *B*(*p*) in an i.i.d. manner as follows: 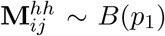 for 1 ≤ *i, j* ≤ *n*_1_ (recurrent connections of layer 1), 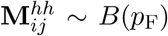 for *n*_1_ + 1 ≤ *i* ≤ *n*_1_ + *n*_2_, 1 ≤ *j* ≤ *n*_1_ (feedforward connections from layer 1 to layer 2),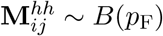 for 1 ≤ *i* ≤ *n*_1_, *n*_1_ +1 ≤ *j* ≤ *n*_1_ +*n*_2_ (feedback connections from layer 2 to layer 1), 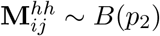 for *n*_1_ + 1 ≤ *i, j* ≤ *n*_1_ + *n*_2_ (recurrent connections of layer 2). Note that in a two-layer network the input is only provided to the first layer and activity is only read out from the second layer. To implement this, we masked both the input layer and the readout layer using binary masks as above to only project input to the first layer and to only read out activity from the second layer, respectively.

To simulate the presence and absence of different types of recurrent connections within a single layer HORN network of *n* nodes, we use the update equation (14) and set the mask matrix **M**^*hh*^ as follows: (i) **M**^*hh*^ = 0 for a network consisting of isolated nodes without self-connections, (ii) 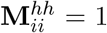 for 1 ≤ *i* ≤ *n* and 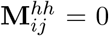 otherwise for a network of isolated nodes with self-connections, (iii) 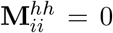 for 1 ≤ *i* ≤ *n* and 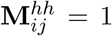 otherwise, resulting in a fully connected network of nodes without self-connections, and (iv) 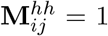 for all *i, j* for a fully connected network of nodes with self-connections. We note that case (iv) is equivalent to that of an unmasked network with the update equation (5).

### Hebbian learning rules

To simulate Hebbian weight updates of the recurrent connection matrix **W**^*hh*^, the weights of the input layer **W**^*ih*^ and the read-out layer **W**^*ho*^ were kept plastic for the BPTT algorithm, but the recurrent weights **W**^*hh*^ were frozen. Instead, a simple additive rule was implemented that modifies the weight of a synaptic connection **W**_*ij*_ from node *j* to node *i* by an additive term

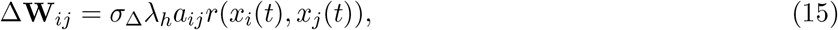

where *σ*_Δ_ ∈ {+1, −1} determines the type of the learning rule (*σ*_Δ_ = +1: Hebb+, *σ*_Δ_ = −1: Hebb-), *λ*_*h*_ denotes the learning rate, *a*_*ij*_ ∈ {0, 1} an activity modulator described below, and *r*(*x*_*i*_(*t*), *x*_*j*_(*t*)) denotes the Pearson correlation between the activity vectors of the nodes *i* and *j* computed over the time steps *t* = *c*, …, *T*, respectively, where *T* = 784 denotes the stimulus length. The parameter *c* controls the starting point in time after which the correlation coefficients are computed. During training, the update rule (15) was applied to the entries of **W**_*ij*_ according to the following logic. For each sample, the distribution of pairwise cross-correlation coefficients *r*(*x*_*i*_(*t*), *x*_*j*_(*t*)) of the evoked HORN activity was calculated for all node pairs. Subsequently, the upper and lower *P* percentiles of this distribution were determined to identify the most strongly correlated and most strongly anticorrelated nodes, excluding self-connections *i, i* (the active connections). The activity modulator *a*_*ij*_ was set to 0 for all node pairs *i, j* for which *r*(*x*_*i*_(*t*), *x*_*j*_(*t*)) did not belong to the set of active connections (the inactive connections). For active connections, the values of *a*_*ij*_ were sampled from a binomial distribution *B*(*p*_*s*_), where *p*_*s*_ is the selection probability. The simulation was performed using the psMNIST data set, see Table S1 for the base HORN parameters. For all HORNs, we disabled the self-connections in the amplitude as well as velocity terms, as these cannot be meaningfully updated with the correlation-based rules, and also disabled bias vectors for the recurrent connections for the same reason. A grid search on *λ*_*h*_ ∈ {5 · 10^−5^, 10^−4^, 5 · 10^−4^, 10^−3^, 5 · 10^−3^}, *c* ∈ {0, 300, 600}, *P* ∈ {1, 5, 10, 50}, *p*_*s*_ ∈ {0.1, 0.5, 1.0} was performed to determine the optimal parameters for each case. For homogeneous HORNs, the following parameters were used for the correlation-based learning rules: Hebb+: *λ*_*h*_ = 10^−4^, *c* = 300, *P* = 5, *p*_*s*_ = 0.5. Hebb-: *λ*_*h*_ = 5 ·10^−4^, *c* = 300, *P* = 5, *p*_*s*_ = 1.0. For heterogeneous HORNs, the following parameters were used: Hebb+: *λ*_*h*_ = 10^−4^, *c* = 300, *P* = 5, *p*_*s*_ = 0.1. Hebb-: *λ*_*h*_ = 10^−4^, *c* = 600, *P* = 5, *p*_*s*_ = 0.1.

### Geometric input

Geometric input was provided to a network by defining non-overlapping receptive fields of size 2 × 2 pixels and covering the 28 × 28 intensity matrix of each MNIST digit by 14 × 14 receptive fields ordered from top left to bottom right of the stimulus matrix in a scanline fashion. Pixel intensities in each of the 196 receptive fields were mapped to the 196 DHO nodes that form a fully connected HORN^*h*^ with node input connection weights and no input bias in a way that preserves geometric relations between the input receptive fields (so that the first DHO node receives input from the top left receptive field, and the last DHO node from the bottom right receptive field). To simulate a flashed stimulus, the network only received input during the first time step. Following the stimulus presentation period, we let the network dynamics unfold for 150 time steps. The whole network was trained using BPTT on the MNIST digit classification task for a readout at time *T* = 150. We trained a HORN^*h*^ network on both the original and a shuffled version of the MNIST digits (with shuffling performed by applying a random but fixed permutation of the 28 × 28 pixel values of each digit).

### Spontaneous activity

For simulating spontaneous activity in HORNs, we trained heterogeneous HORN networks of size 16 on sMNIST (see Table S.1 for parameter values). We used random initial conditions for each DHO node with both the initial amplitudes *x* and velocities 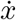 for *t* = 0 sampled from a Gaussian distribution *N* (0, 30). To model spontaneous activity, the DHO nodes in the network were subjected to random instantaneous deflections to their velocity 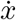 in the form of delta pulses induced by i.i.d. Poisson processes with a rate of *λ* = 0.005 and an amplitude Of 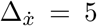. To model the transition into and out of a stimulus-driven regime, we simulated the network dynamics in three periods. In the first periods, the network was spontaneously active for a period of 1000 time steps, after settling the network activity for 500 time steps. In the second periods lasting 784 timesteps, spontaneous activity was switched off, and an sMNIST stimulus was presented. The third period lasting 1000 time steps represents the time after stimulus offset when the stimulus-dependent input was replaced again by spontaneous activity. PCA analysis was performed on windowed network activity (window length 20 time steps). In particular, windows of spontaneous activity were sampled starting at *t* = 500, and windows of evoked activity starting at *t* = 1764 (i.e. the windows spanning 20 time steps prior to the readout time). PCA was performed using the implementation in the Python package scikit-learn (sklearn.decomposition.PCA).

## Data and materials availability

Model implementations and simulation scripts will be made publicly available upon publication. All data sets used in this study are publicly available.

The authors thank Martin Vinck, Andreas Bahmer, and Andreea Lazar for their helpful discussions and comments on an earlier version of this manuscript. We thank Claudia Kernberger for her help with some of the illustrations. We acknowledge computing support from the Ernst Strüngmann Institute and the Max Planck Computing and Data Facility.

