## Supplementary Materials for "The functional role of oscillatory dynamics in neocortical circuits: a computational perspective"

### Supplementary Text

#### 1 Relation to the Wilson-Cowan model

The Wilson-Cowan (WC) equations [1] are a mean-field model that describes the dynamics of two recurrently coupled populations of neurons, an excitatory (E) and an inhibitory (I) one. Denoting by $E(t)$ and $I(t)$ the instantaneous firing rate of the excitatory and inhibitory population, respectively, the Wilson-Cowan equations are

$$\begin{aligned}\tau\dot{E}(t) &= -E(t) + (1 - rE(t))f_E\left[w_{EE}E(t) - w_{EI}I(t) + h_E(t)\right] \\ \tau\dot{I}(t) &= -I(t) + (1 - rI(t))f_I\left[w_{IE}E - w_{II}I + h_I(t)\right],\end{aligned} \tag{S1}$$

where $\tau$ is the characteristic time scale of the system, $w_{XY}$ is the coupling strength of the subpopulation $X$ to $Y$ (with $X, Y \in \{E, I\}$), $h_X$ is the external input of the subpopulation $X$, and $f_X$ is a non-linear (usually sigmoidal) activation function of the subpopulation $X$. Here, we use a slightly modified version of the Wilson-Cowan model, as in [2], which is known to express oscillations. Here, the activation function $f_X$ is given by

$$f_X = \frac{1}{1 + \exp(-a_X(X - \theta_X))}, \tag{S2}$$

where $\theta_X$ define the center and $a_X$ the steepness of the nonlinearity for the population $X = E, I$, respectively.

Following [2], we take the external inhibitory and excitatory inputs $h_I$ and $h_E$ as free parameters and fix the values of the other parameters as $w_{EE} = 13, w_{EI} = 6, w_{IE} = 12, w_{II} = 3, \tau = 1, a_e = 1.3, a_i = 2, \theta_e = 4, \theta_i = 1.5$ for our analysis.

In the $h_E$-$h_I$ subspace, we obtain a bifurcation diagram (fig. S21) that partitions the state space of the system into regions with three dynamically distinct behaviors: (i) asymptotically decaying dynamics, (ii) oscillatory decaying dynamics, and (iii) dynamics expressing sustained oscillations (or limit cycles). The system trajectories in each subspace behave qualitatively in the same way as the ones produced by a damped harmonic oscillator subject to different parameter and input configurations. The dynamics in region (i) are equivalent to the ones produced by a supercritically damped DHO, the ones of region (ii) correspond to a subcritically damped DHO, and the dynamics in region (iii) correspond to the case of a driven subcritically damped DHO (or a node with velocity feedback; see Section 2.1).

The WC model allows for a rich repertoire of qualitatively different dynamics. However, it is difficult to tune its oscillatory properties, and in particular the frequencies of the oscillations (fig. S21), given that the model has 11 parameters. DHOs on the other hand have two easily interpretable parameters $\omega$ and $\gamma$, describing their oscillation frequency and decay behavior, respectively (fig. S1 A and B). Despite its simplicity, the DHO model (fig. S1) is able to qualitatively reproduce the different dynamics expressed by the WC model for the parameter values considered above (fig. S21).

#### 2 Dynamics of DHO nodes

In HORNs, one DHO node models the aggregate activity of a cortical microcircuit such as a (P)ING circuit or a cortical column, consisting of both an excitatory and an inhibitory population of neurons. Local recurrent connectivity within each microcircuit results in feedback input of the circuit to itself (see Section 1). To model this feedback activity, we introduce two forms of feedback-connectivity for each DHO node: a velocity and an amplitude feedback connection. Here, we analyze the impact of each type of feedback on the dynamics of a node, as well as their combined influence. We start by revisiting the general expression of our DHO node dynamics

$$\ddot{x}(t) + 2\gamma\dot{x}(t) + \omega^2 x(t) = \alpha\sigma\left(I\left(t\right) + F\left(x\left(t\right), \dot{x}\left(t\right)\right)\right), \tag{S3}$$

where $x(t)$ denotes the oscillator's amplitude, $\gamma > 0$ the damping factor, $\omega > 0$ the natural angular frequency, and $\alpha > 0$ an excitability parameter. $\sigma(\cdot)$ denotes an input nonlinearity that models local resource constraints and is chosen as $\tanh(\cdot)$ unless otherwise stated. $I(t)$ denotes a time-dependent external input. In the first part of our analysis, we will only work with free oscillations, setting $I(t) = 0$. The feedback input of the node to itself is $F(x(t), \dot{x}(t)) = vx(t) + w\dot{x}(t)$, $(v, w \in \mathbb{R})$. Combining the above, the DHO equation can be written as

$$\ddot{x}(t) + 2\gamma\dot{x}(t) + \omega^2 x(t) = \alpha \tanh\left(vx + wx\right). \tag{S4}$$

#### 2.1 Velocity-feedback DHO

Let us first considering only the velocity feedback in isolation, setting $v = 0$ in (S4). This ODE can be rewritten as a two-dimensional system:

$$\begin{aligned} \dot{x} &= y \\ \dot{y} &= -2\gamma y - \omega^2 x + \alpha \tanh\left(wy\right). \end{aligned} \tag{S5}$$

The eigenvalues of the Jacobian of this system are are given by

$$\lambda_{1,2} = \frac{\alpha w - 2\gamma \pm \sqrt{\left(\alpha w - 2\gamma\right)^2 - 4\omega^2}}{2} \tag{S6}$$

and cross the imaginary axis at the value $w_c = 2\gamma/\alpha$, which characterizes a Hopf bifurcation (fig. S22) [3]. In terms of system dynamics, this means that the node starts to express sustained oscillations if the velocity self-connection term $w$ is greater than $w_c$.

#### 2.2 Amplitude-feedback DHO

To consider amplitude feedback in isolation, we set $w = 0$ in (S4) and get

$$\ddot{x}(t) + 2\gamma\dot{x}(t) + \omega^2 x = \alpha \tanh(vx). \tag{S7}$$

Rewriting the second-order differential equation (S7) as a system of first order ODEs

$$\begin{aligned} \dot{x} &= y \\ \dot{y} &= -2\gamma y - \omega^2 x + \alpha \tanh\left(vx\right), \end{aligned} \tag{S8}$$

we can calculate the nullclines of this system as

$$\dot{x} = 0 \Rightarrow y = 0, \tag{S9}$$

$$\dot{y} = 0 \Rightarrow y = \frac{\alpha \tanh\left(vx\right) - \omega^2 x}{2\gamma}. \tag{S10}$$

The intersections of the nullclines (fig. S23) define the fixed points of the system that we denote by $(x^*, y^*)$. As all fixed points of the system satisfy $y^* = 0$, we will refer to the fixed point $(x^*, 0)$ as $x^*$ from now on, unless otherwise stated.

Combining (S9) and (S10), we obtain the following transcendental equation

$$\tanh vx^* = \frac{\omega^2}{\alpha} x^*, \tag{S11}$$

which cannot be solved analytically. To obtain solutions, we compute the intersection between the nullclines numerically (fig. S23). This allows us to study the behavior of the fixed points of the system $x^*$ for different values of $v$ (fig. S24).

The bifurcation diagram (fig. S24) shows that the nodes undergo a pitchfork bifurcation at the critical point $v_c = \frac{\omega^2}{\alpha}$. This means that our nodes have different stationary behaviors for different intervals of the values of $v$: (i) Subcritical case: The node has a single stable solution at (0,0) when $v \leq \frac{\omega^2}{\alpha}$. (ii) Supercritical case: Once we cross the bifurcation point $v = v_c$, the origin loses stability and two new stable solutions arise, given by (S11). In the latter case, the stationary solution depends on the initial conditions, see (fig. S25).

Although the solutions of (S11) cannot be analytically obtained, we can calculate the bounds to the solutions. Consider $x' = x^* \frac{\omega^2}{\alpha}$. We can then rewrite (S11) as

$$\tanh v \frac{\alpha}{\omega^2} x' = x'. \tag{S12}$$

When $v \to \infty$ we have $x' \to \pm 1$, $x^* = \pm \frac{\alpha}{\omega^2}$ as the asymptotic values for the fixed points (fig. S23 black dots, fig. S24 dashed lines).

##### 2.3 Amplitude and velocity feedback - Symmetric Bogdanov-Takens Bifurcation

The ODE for a DHO node subject to both velocity and amplitude feedback is

$$\ddot{x}(t) + 2\gamma \dot{x}(t) + \omega^2 x(t) = \alpha \tanh\left(vx(t) + w\dot{x}(t))\right), \tag{S13}$$

where $v$ and $w$ denote the amplitude and velocity feedback parameter, respectively.

The complete system of feedbacks has the normal form of a symmetric Bogdanov-Takens bifurcation, with bifurcations in the parameter space $(v, w)$ as depicted in (fig. S3); for a detailed discussion, see [4]. We will briefly break down this codimension-two bifurcation by describing its most important bifurcation curves: at $v = 0$ the system undergoes a pitchfork bifurcation. At $w = 0$, for $v < 0$ and at $v = w$ (for $v > 0$) the system undergoes Hopf bifurcations. For $v > w > 0$, more complicated global bifurcations occur due to collisions between the limit cycles of the trivial fixed point and the non-trivial ones (fig. S3).

#### 3 Driven DHO node

We now consider the model dynamics in the driven regime, i.e. $I(t) \neq 0$ ( S4).

##### 3.1 Constant Input and amplitude feedback - Cusp Bifurcation

Here, we consider the case of a constant input, $I(t) = b$, and its interaction with amplitude feedback. Note that the constant input is given by the bias term of a given node. This results in an equation for the fixed points which is similar to the one of the undriven undriven case (S11)

$$\alpha \tanh\left(vx^* + b\right) = \omega^2 x^*. \tag{S14}$$

Solving for $v$, we obtain

$$v = \frac{\tanh^{-1}\left(\frac{\omega^2}{\alpha} x^*\right) - b}{x^*}. \tag{S15}$$

The parameter $b$ breaks the symmetry of (S15) (fig. S26, top). Given a fixed $b \neq 0$, the system undergoes a saddle-node bifurcation as $v$ passes $x_m$, the local minimum of (S15). We can calculate $x_m$, the local minimum and $x_i$, the inflection point, by setting $\partial v / \partial x^* = 0$ and obtain

$$b + \frac{\frac{\omega^2}{\alpha} x_m}{1 - \left(\frac{\omega^2}{\alpha} x_m\right)^2} - \tanh^{-1}\left(\frac{\omega^2}{\alpha} x_m\right) = 0 \tag{S16}$$

and

$$b - \frac{\frac{\omega^2}{\alpha} x_i}{1 - \left(\frac{\omega^2}{\alpha} x_i\right)^2} + \tanh^{-1}\left(\frac{\omega^2}{\alpha} x_i\right) = 0 \tag{S17}$$

as the two solutions. This symmetry breaking creates two branches on which the fixed points of the system lie. One branch, which is stable, persists for all values of $b$, and the other branch, as mentioned above, is a saddle. To obtain the bifurcation point, we sequentially solve equations (S16) and (S15) numerically. Equation (S16) allows us to compute the local minimum $x^* = x_m$, which we insert into (S15) to obtain $v_m = v(x_m)$. The saddle-node bifurcation point is then defined by the tuple $(x_m, v_m)$. In (fig. S26) we see that $b$ influences the position of the saddle-node bifurcation. To understand how $b$ can affect the location of the bifurcation point $(x_m, v_m)$, we now fix $v$ and consider the subspace $(b, x^*)$. We find that in the limits $v \to \pm\infty$ the fixed points converge to $x^* \to \pm\frac{\alpha}{\omega^2}$, respectively (fig. S27). We find that for $v < \frac{\omega^2}{\alpha}$ (subcritical node), the position of the fixed point is sensitive to the external current amplitude $b$ (fig. S27). For $v > \frac{\omega^2}{\alpha}$ (supercritical node), we find that the regions with three fixed points are bounded by the local minima and maxima of $b(x^*)$. We can obtain this range by setting $\partial b / \partial x^* = 0$ in (S15) and get

$$x_{\text{min,max}} = \pm \frac{\sqrt{v - \frac{\omega^2}{\alpha}}}{\frac{\omega^2}{\alpha}\sqrt{v}}. \tag{S18}$$

Plugging these values into (S15), we get

$$b_{\pm} = \tanh^{-1}\left(\pm\frac{\sqrt{v - \frac{\omega^2}{\alpha}}}{\sqrt{v}}\right) \mp \frac{\sqrt{v}\sqrt{v - \frac{\omega^2}{\alpha}}}{\frac{\omega^2}{\alpha}}, \tag{S19}$$

for the amplitude interval $[b_-, b_+]$ for which there exist three fixed points (fig. S28, red lines).

The complete bifurcation diagram in the parameter subspace $(v, b)$ is obtained by concatenating the two previous bifurcation diagrams. The resulting bifurcation diagram (fig. S28) describes a cusp bifurcation, where the boundaries of the diagram are given by $b_-(v)$ and $b_+(v)$ (see (S19)).

##### 3.2 Constant input and Velocity-feedback

Considering only the velocity feedback by setting $v = 0$ in (S4), we obtain

$$\begin{aligned} \dot{x} &= y \\ \dot{y} &= -2\gamma y - \omega^2 x + \alpha \tanh(wy + b). \end{aligned} \tag{S20}$$

This system has the fixed point

$$\begin{aligned} y^* &= 0 \\ x^* &= \frac{\alpha \tanh(b)}{\omega^2}. \end{aligned} \tag{S21}$$

The trace and determinant of the Jacobian $J$ at the fixed point are

$$\begin{aligned} \operatorname{tr} J &= -2\gamma + \alpha w \operatorname{sech}^2(b), \\ \det J &= \omega^2 \end{aligned} \tag{S22}$$

and cross the imaginary axis at the curve

$$w_c(b) = \frac{2\gamma}{\alpha \operatorname{sech}^2(b)}, \tag{S23}$$

which characterizes a Hopf bifurcation (fig. S31) [3]. In terms of system dynamics, this means that the node starts to express sustained oscillations if the velocity self-connection term $w$ is greater than $w_c$.

##### 3.3 Harmonic Input

Given the oscillatory nature of the recurrent input that each node in a HORN receives, we study here how a single node subject to amplitude feedback responds to harmonic stimuli, to better understand its dynamics. Consider as input $I(t) = A\sin(\omega_i t)$, where $\omega_i$ denotes the input frequency. Inserting this input into (S7), we obtain the following nonautonomous equation

$$\ddot{x} + 2\gamma\dot{x} + \omega^2 x = \alpha \tanh\left(vx + A\sin(\omega_i t)\right). \tag{S24}$$

The parameter space is now composed of the damping ratio $\gamma$, the natural frequency $\omega$, the excitability $\alpha$, the amplitude feedback $v$, the amplitude of the input $A$ and the input frequency $\omega_i$.

Here, we study the node as an autonomous frozen system, where the dynamics of the nodes (fig. S26 bottom) are ruled by fixed points that change their positions due to the harmonic input. We will call these moving fixed points, also known as moving stable state [5]. One way to picture this scenario is to consider that our bifurcation diagram is periodically and continuously transitioning between the upper panels of (fig. S26). In this scenario, we can identify three different regions: In region 1, the system has a single moving fixed point (fig. S26, red). In region 2, two alternating saddle-node bifurcations dynamically create a continuous 3 to 1 fixed points regime (fig. S26, blue). The detachment of a stable branch due to its collapse to the unstable branch, that is, the saddle-node bifurcation, is known as a type of bifurcation-induced tipping (B-tipping) [6], where the parameter change path crosses a bifurcation. In region 3, the system has 3 nonvanishing moving fixed points (fig. S26 green). The fixed point oscillating around the origin is unstable, whereas the other two are stable branches. Here, system dynamics can be captured by the single stable branch or it is juggled between the two stable branches. The three regions are delimited by the non-driven bifurcation point $v = \frac{\omega^2}{\alpha}$ (fig. S26, dashed dot line) and the position of the local minima $v(x_m(A))$ (see (S16) with $b = A$ and fig. S26, dashed line). This analysis allows us to gain a better understanding of the DHO node dynamics for harmonic inputs.

We proceed with a brief computational approach, to give an overview of the dynamics of the system for different initial conditions. Given the nonautonomous nature of the ODE, the initial conditions of the system are defined by the triple $(x_0, y_0, t_0)$, where $t$ denotes time. The absence of dependence of stationary behavior on initial conditions can result in two different types of attractiveness in nonautonomous systems, namely, forward and pullback attractiveness [7]. For the DHO nodes in our networks, such a dependency could imply two different modes of computation. In the first case, the computation relies on the initial state of the node. In the other case, the computation exploits the stationary state of the node.

Using simulated single node dynamics, we numerically investigate the types of attractiveness of the DHO nodes for different parameter values and initial conditions. Since our input is harmonic, any initial condition $(x_0, y_0, t_0)$ can be expressed as $(x_0, y_0, t_0 + k\frac{2\pi}{\omega_i})$, with $k \in \mathbb{Z}$ due to the periodicity of the input. Without loss of generality, we restrict our analysis to the time window $[0, \frac{2\pi}{\omega_i}]$.

We start by inspecting pullback attractiveness. From (fig. S29) we observe that subcritical and critical nodes seem to display pullback attraction, but for supercritical nodes this does not always seem to be the case.

To investigate forward attractiveness, we inspect the influence of different initial positions $x_0$ on system dynamics. From (fig. S30) we conclude that subcritical and critical nodes seem to display forward attraction, while again this does not always seem to be the case for supercritical nodes.

In this section, we analyzed DHO node dynamics resulting from harmonic input and found that node dynamics did not just depend on parameter values but also on on initial conditions, including phase difference with the input.

### 4 Resonance

Here, we inspect how DHO nodes with feedback connections resonate to harmonic input of the form $I(t) = \sin(\omega_i t)$. Each DHO node performs a gain modulation of its input depending on its natural frequency $\omega$ and the input frequency $\omega_i$ (S7). The amplitude feedback parameter $v$ influences the peak position of the gain curves (Fig. S4). Moreover, the initial plateau $G = 1$ becomes less than one for negative values of $v$, indicating that for negative values of amplitude-feedback, the DHO nodes becomes a band-pass filter instead of low-pass filter in the case of $v = 0$ (fig. S4). We could not find any influence the velocity-feedback term $w$ on the gain curve in our simulations (data not shown).

#### Supplementary Figures

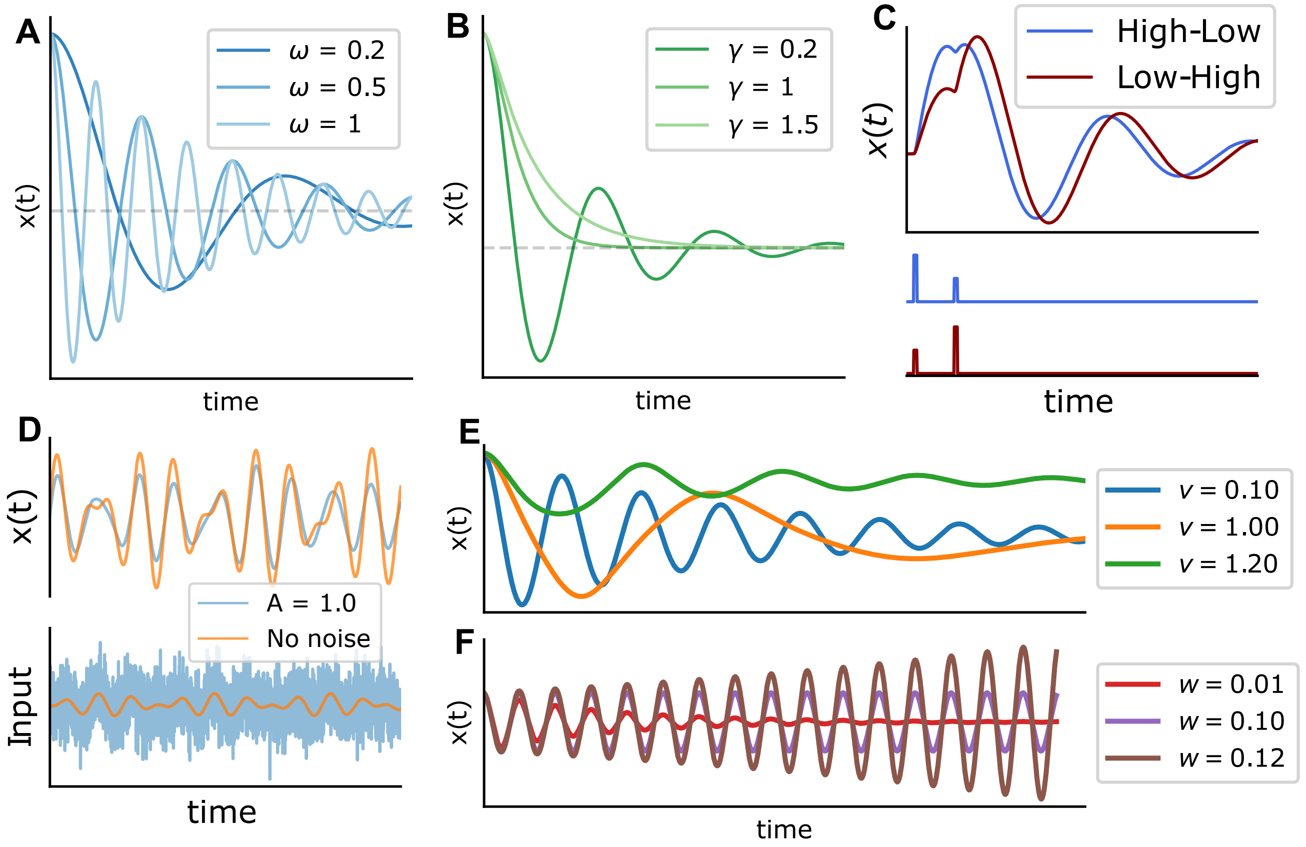


**Fig. S1**. Relaxation dynamics of a single DHO node ($\omega = 1$, $\gamma = 0.05$, $\alpha = 1$). A. Effect of the natural frequency parameter $\omega$. B. Effect of the damping parameter $\gamma$. C. Sequence sensitivity of a DHO. The sequence order of two delta pulses with different amplitudes gets encoded in the oscillation phase. D. DHO node response (top) to two different input signals (bottom). Input signals ans response color coded. Orange: input without noise, blue: input with additive white Gaussian noise. Note the similar amplitude trajectory $x(t)$ in both cases. E. Effect of the amplitude feedback parameter $v$ on the relaxation dynamics of a DHO node (initial condition $(x_0, y_0) = (1, 0)$). The critical value of $v$ is $v_c = 1$. Note the emergence of two stable attractors for $v > v_c$. Relaxation dynamics to one of the attractors shown (green line). For $v < v_c$, only one attractor exists. F. Effect of the velocity feedback parameter $w$ on the relaxation dynamics of a DHO node (initial condition as in E). The critical value for $w$ is $w_c = 0.1$. Note the emergence of a limit cycle for $w > w_c$ (Hopf bifurcation).

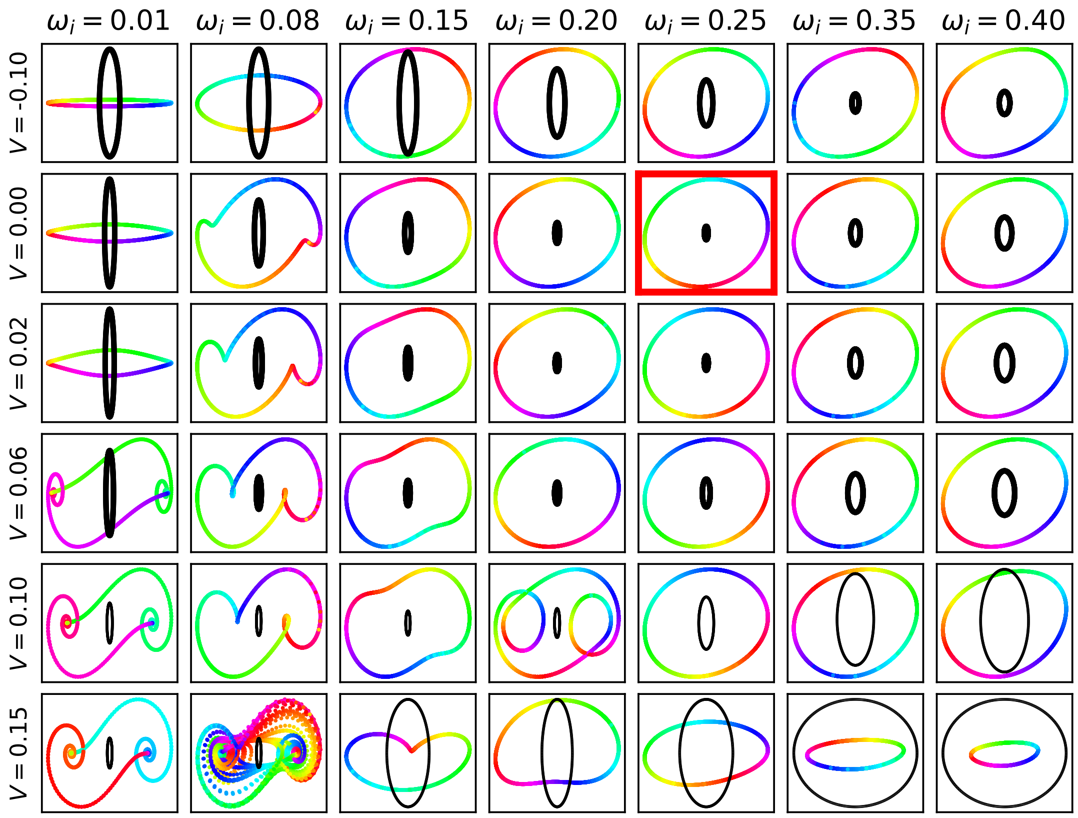


**Fig. S2**. Dynamics of a single DHO node with $\omega = 0.25$, $\gamma = 0.05$, $\alpha = 1$ driven by a harmonic input with an amplitude of $A = 1$ for different input frequencies $\omega_i$ and self-connection strengths $V$. Each panel shows the phase space of the stationary dynamics of the DHO node (colored curve, color-coded according to the input phase), due to the driving harmonic input (black curve, node circle). The panel corresponding to $\omega_i = 0.25$, $v = 0$ (red frame) shows the case of a simple damped harmonic oscillator without feedback.

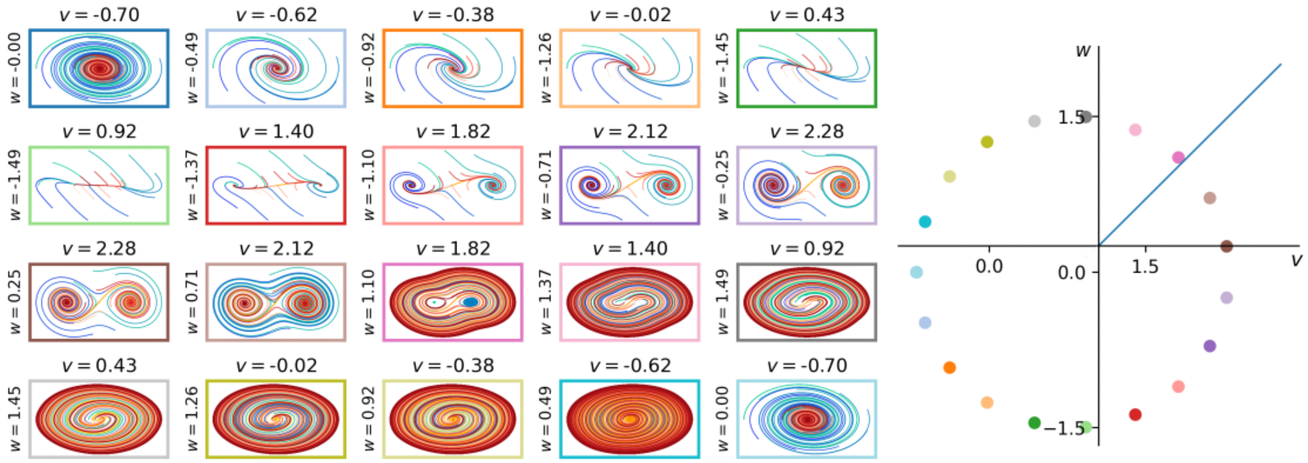


**Fig. S3**. Bifurcation diagram of a DHO node with amplitude and velocity feedback. Left: Examples of phase portraits for nodes with different combinations of parameters $v, w$. Cold colors: initial conditions of an outer circle. Warm colors: initial conditions of an inner circle. The frame colors indicate the position on the bifurcation diagram (right). Right: Qualitative bifurcation diagram for a DHO node in $(v, w)$ space, origin centered at $(v_c, w_c)$ for fixed parameters values $\omega = 1, \alpha = 1, \gamma = 0.05$. Blue blue line indicates the identity $w = v$ (with $v > v_c$ and $w > w_c$). Colors of points match frame colors on the left.

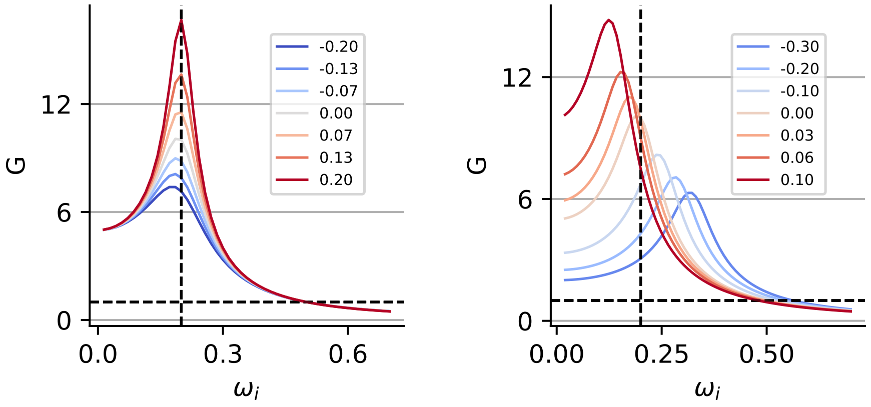


**Fig. S4**. Dependence of the gain function $G(\omega_i)$ as intrinsic receptive fields of a DHO node ($\gamma = 0.05$, $\omega = 0.2$, $\alpha = 0.2$) calculated using sinusoidal input of varying frequency $\omega_i$ and unit amplitude on feedback parameter values. Left: Dependence of $G(\omega_i)$ on velocity feedback parameter $w$ (diagonal entry in coupling matrix), values color coded. Right: Dependence of $G(\omega_i)$ on amplitude feedback parameter $v$, values color coded.

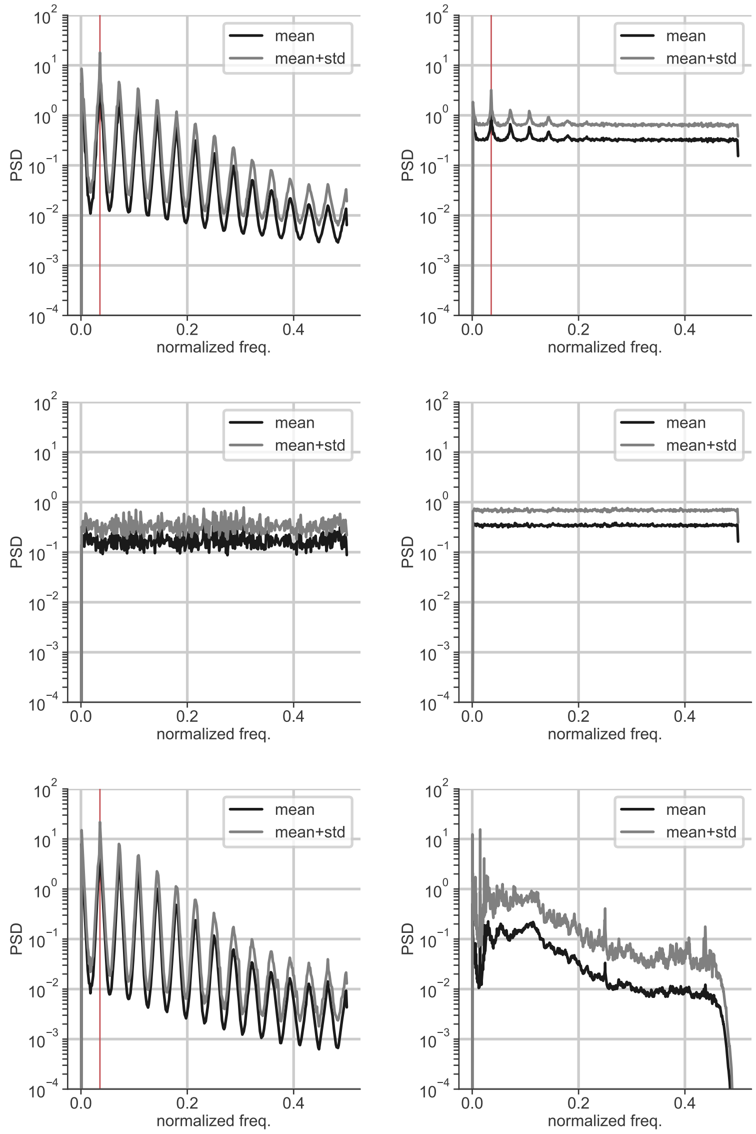


**Fig. S5**. Power spectral densities for groups of samples from different data sets. Plots show mean and std.-dev. in PSD computed over 1000 samples (`scipy.periodogram` with `fs=1.0`). Vertical axes show PSD, horizontal axes show normalized frequency $f$. The red line indicates the line frequency of MNIST ($2\pi/28$). From top left to bottom right: sMNIST, sMNIST corrupted by $N(0, 1)$ additive white Gaussian noise, psMNIST, psMNIST corrupted by $N(0, 1)$ additive white Gaussian noise, EMNIST, Spoken Digits Data Set.

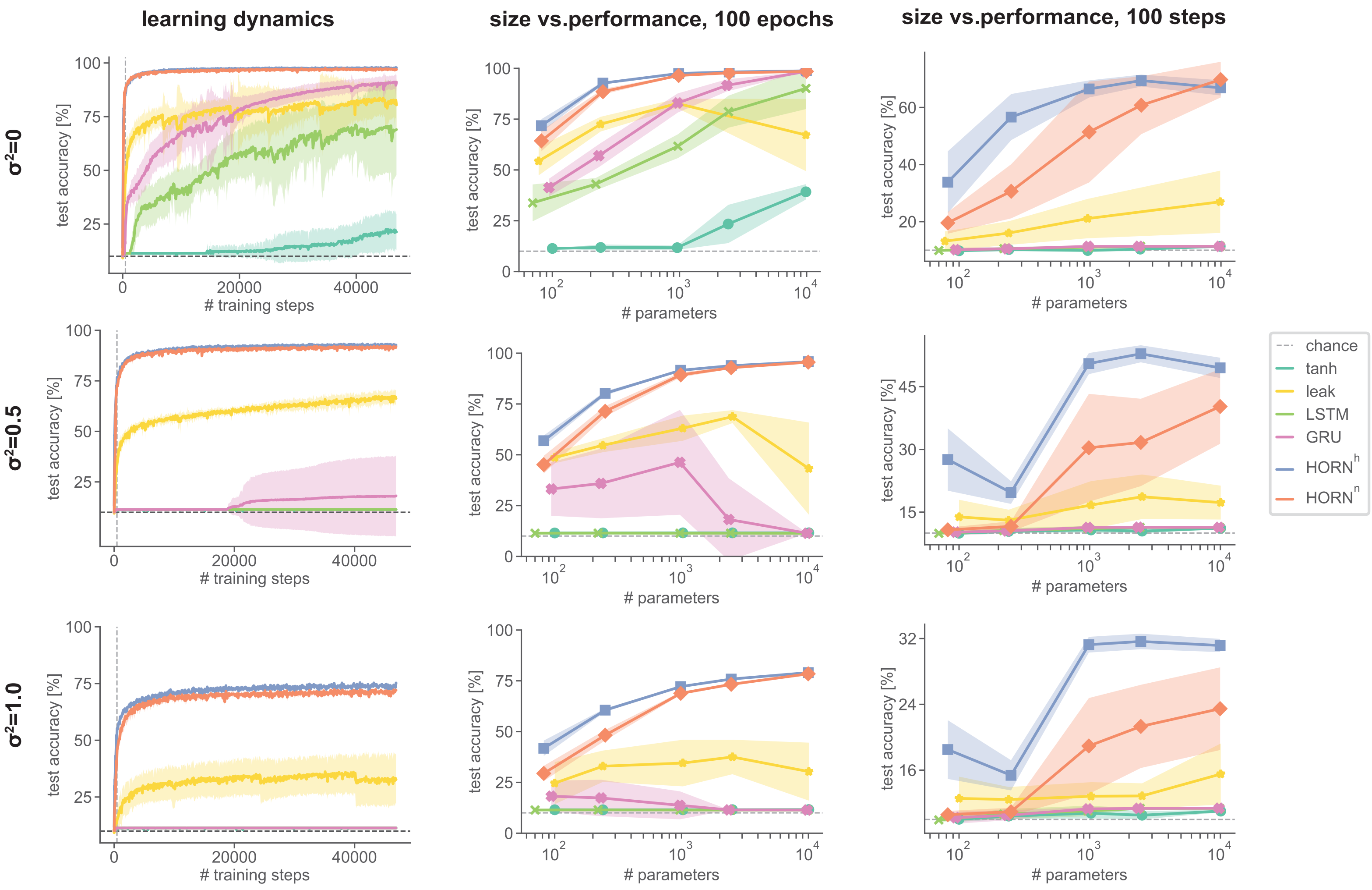


**Fig. S6**. Learning dynamics and performance for different RNN architectures on sMNIST. Left column: Learning dynamics for a network with 2500 trainable parameters. Middle and right columns: Maximal accuracy after 100 learning epochs and after 100 BPTT update steps as a function of the number of trainable parameters, respectively. The rows show different noise levels $\sigma^2$. Lines show mean over 10 network instances with random weight initialization, shaded areas standard deviation. Vertical dashed lines mark the end of the first learning epoch.

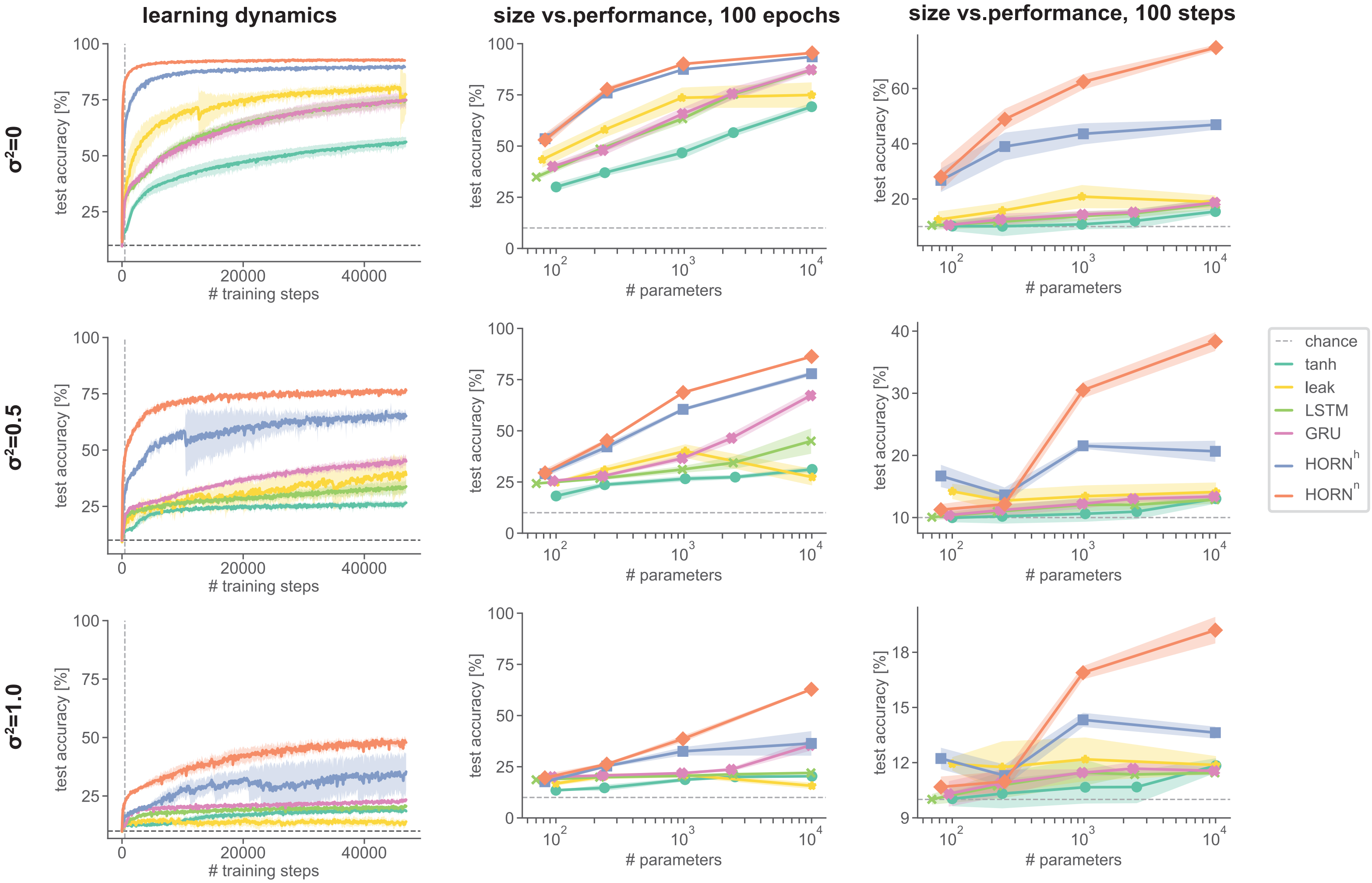


**Fig. S7**. Learning dynamics and performance for different RNN architectures on psMNIST. Panels as in Fig. S6.

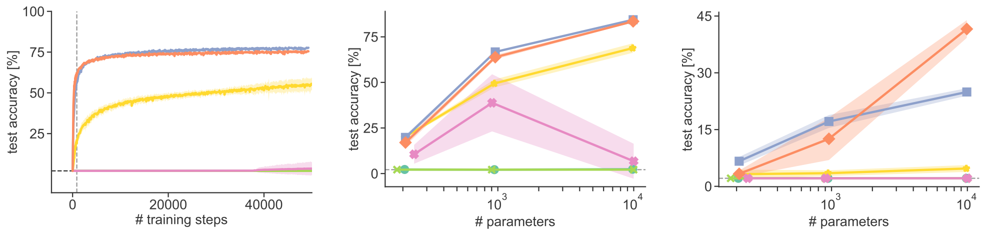


**Fig. S8**. Performance of different RNN architectures on serialized EMNIST (balanced) data set. Left column: Learning dynamics for a network with 2500 trainable parameters. Middle and right columns: Maximal accuracy after 100 learning epochs and after 100 BPTT update steps as a function of the number of trainable parameters, respectively. Lines show mean over 10 network instances with random weight initialization, shaded areas standard deviation. Colors as in fig S6.

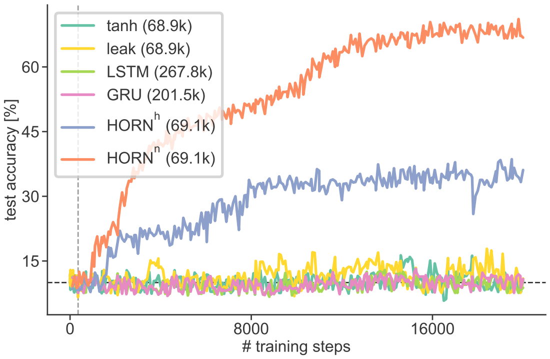


**Fig. S9**. Classification accuracy as a function of training steps for different RNN architectures when trained on SDDS for 50 epochs (batch size 64). All networks had 256 nodes (number of trainable parameters in parentheses). HORN parameters as in Table S1.

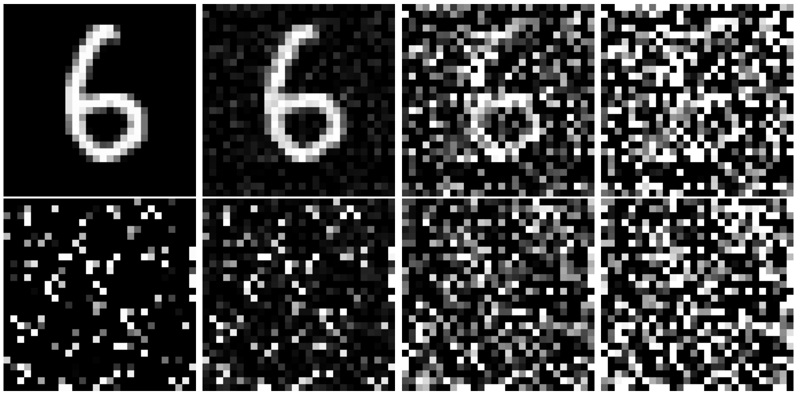

**Fig. S10**. Example digit from MNIST data set in unshuffled (top row) and shuffled form (bottom row). Columns show different levels of additive white Gaussian noise applied per pixel with $\sigma^2 = 0, 0.1, 0.5, 1.0$ (from left to right). Stimuli are clipped to values in $[0, 1]$ after noise application.

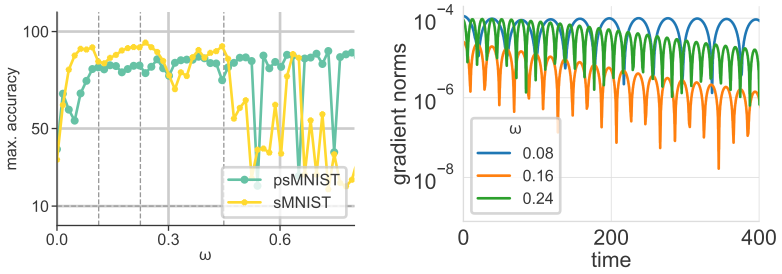


**Fig. S11**. Influence of the natural frequency parameter $\omega$ on performance and gradient propagation in homogeneous HORN network (64 units, $h = 1$, $\gamma = 0.01$, $\alpha = 0.04$, no self-connection term $v$) for sMNIST and psMNIST data sets. Left: Maximum test accuracy attained in 20 training epochs as a function of $\omega$. Vertical dashed lines indicate the fundamental frequency of the sMNIST data set $2\pi/28$ and its harmonics. Note the stronger dependence of the network performance on $\omega$ for sMNIST, but not for psMNIST. Right: Oscillating $L^\infty$ gradient norms in an untrained network as a function of time on the sMNIST test set. Lines show mean $||\frac{\partial L}{\partial x_i}||_\infty$ averaged over all samples of the test set. Note the dependence of the frequency of the gradient oscillations on $\omega$.

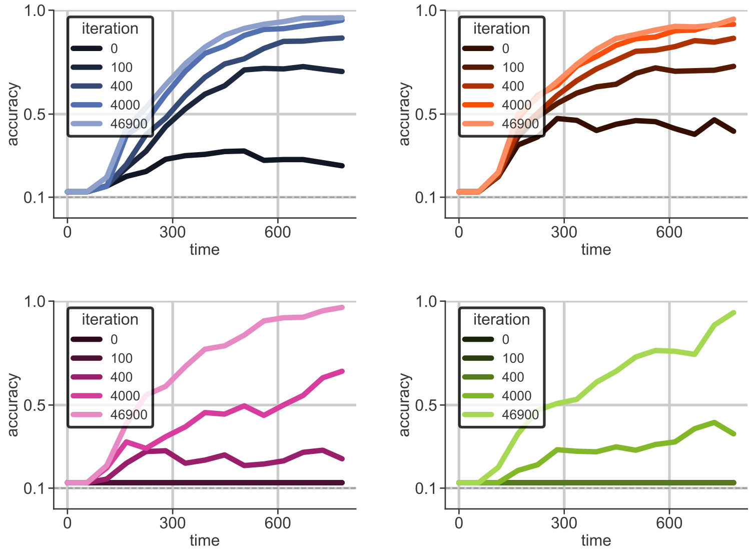


**Fig. S12**. Dynamics of stimulus decodability by linear SVM from RNN activity vector for different architectures for different network instances during training on sMNIST for 100 epochs, from top left to bottom right HORN$^h$, HORN$^n$, LSTM, GRU. Color lightness indicates number of BPTT training iterations.

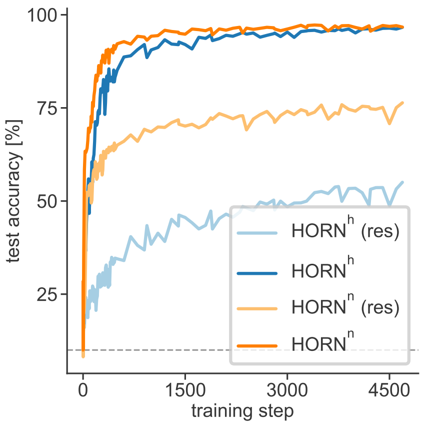


**Fig. S13**. Learning dynamics of full and reservoir HORNs of size 128 during 10 training epochs on sMNIST. For reservoir HORNs, only input and readout layers are trained with BPTT.

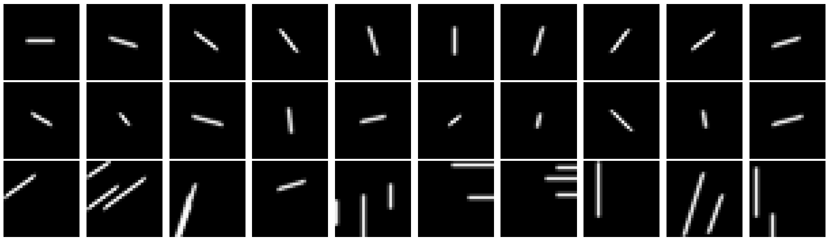

**Fig. S14**. Samples of Line Segments data set. Top row: Angles present in LSDS(10,1,10,10,c). Middle row: Samples from LSDS(32,5,11,c) (referred to as LSDSa). Bottom row: Samples from LSDS(10,3,8,24,r) (referrred to as LSDSb).

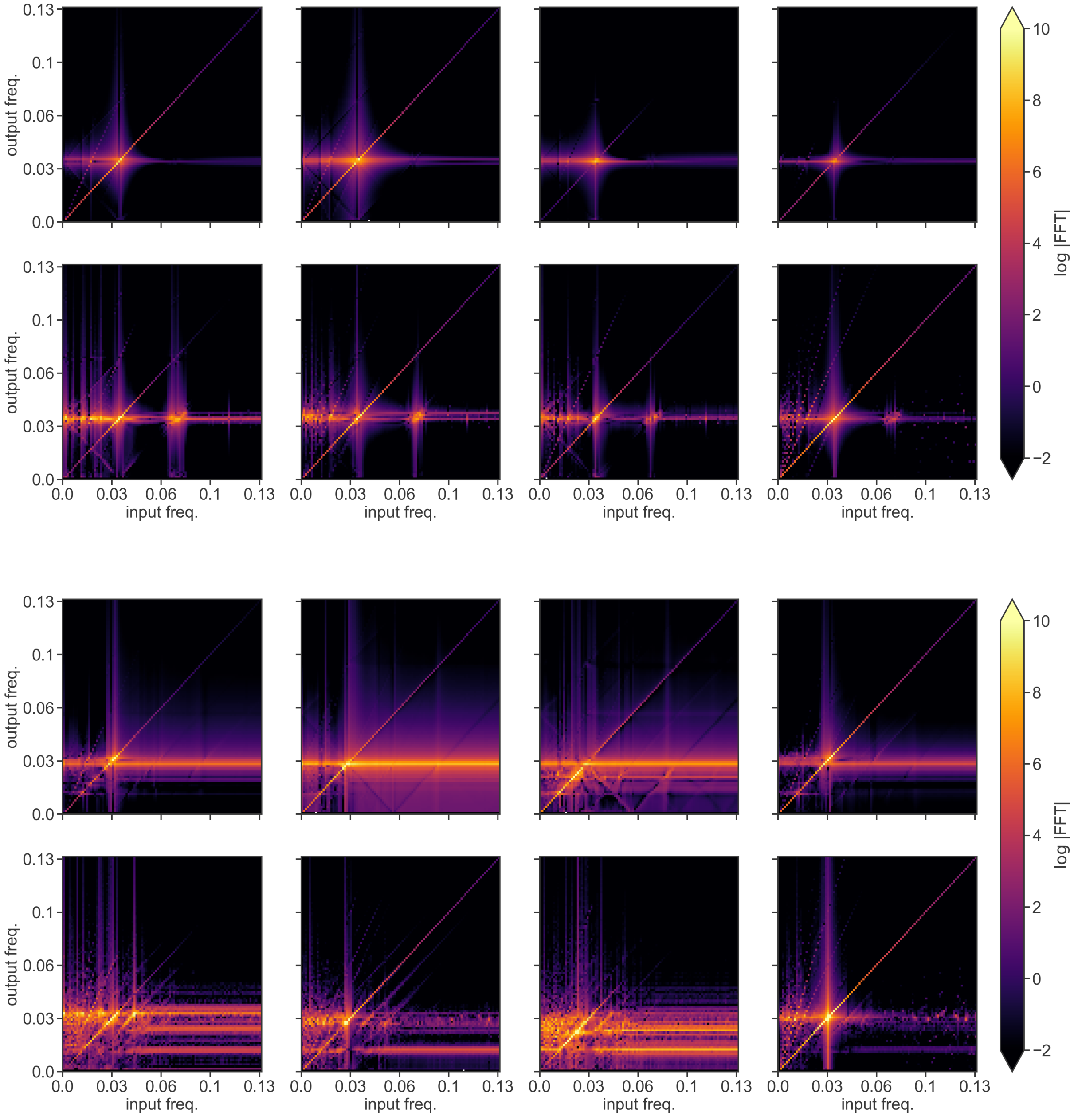


**Fig. S15**. Resonance behavior of DHO nodes in HORN networks (16 nodes) in response to sinusoidal input stimulation before and after training on sMNIST for 100 epochs. Top panel shows homogeneous HORN, bottom panel shows heterogeneous HORN. Each panel shows four nodes from the network in an untrained state (first row) and after training (second row). Each matrix shows the logarithm of the power spectral density (PSD) of a node's activity across frequency bands (output freq., vertical axis) as a function of the input frequency of the sinusoidal stimulus (input freq., horizontal axis). Color scale on the right, PSD values clipped to a range between $10^{-2}$ and $10^{10}$ to enhance visibility. Note that the bright horizontal lines correspond to the natural frequencies of the nodes, vertical lines indicate resonance at input frequencies around the nodes natural frequency, and bright diagonal lines indicate entrainment. Nodes in HORNs develop complex resonance behavior (bottom rows in both panels).

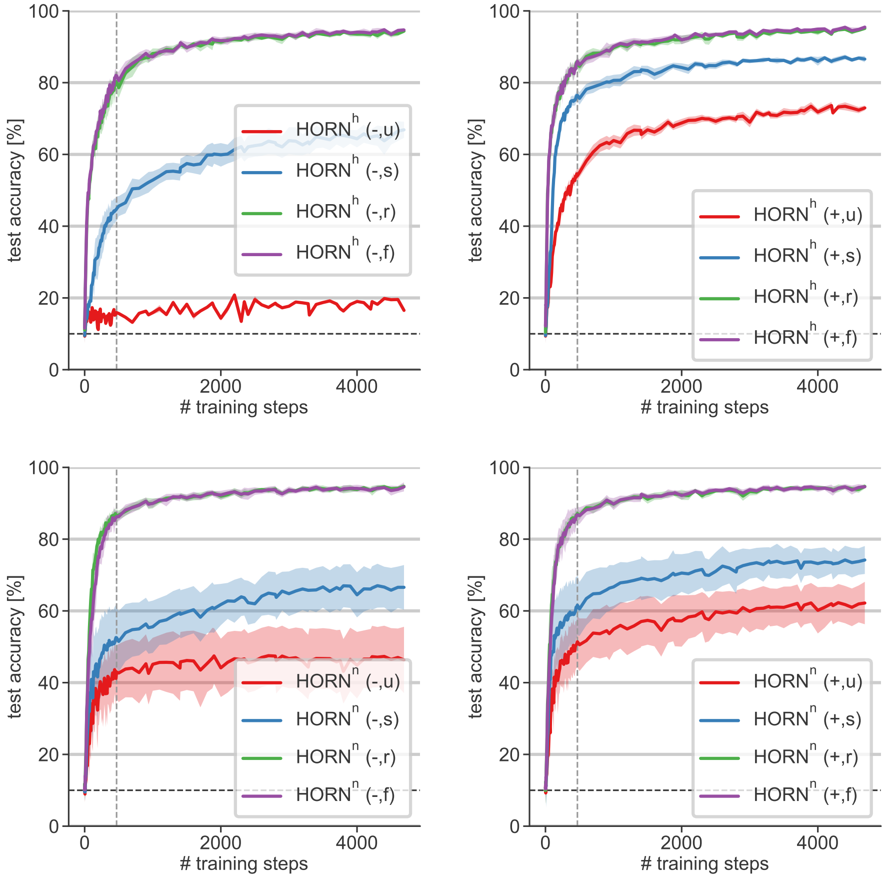


**Fig. S16**. Learning dynamics of HORN networks (32 nodes) trained on sMNIST for 10 epochs under different connectivity constraints. Lines show mean for 10 random weight initializations, shaded areas standard deviation. Network label in parentheses indicates presence of feedback parameters and network connectivity pattern: $-$ $(+)$ indicates absence (presence) of amplitude feedback connection, "u" fully unconnected network of isolated nodes ($\mathbf{W}^{hh} = 0$), "s" indicates network of isolated nodes with self-connections in $y$ state variable ($\mathbf{W}^{hh} = \mathrm{diag}(w_{11}, \ldots, w_{nn})$), "r" indicates fully connected network without velocity self-connections ($\mathbf{W}^{hh}_{ii} = 0$), "f" indicates fully connected network with velocity self-connections. Top row: homogeneous HORN, bottom row: hetergeneous HORN. Left column: no apmlitude feedback-connections in $x$, right column: amplitude feedback connections present.

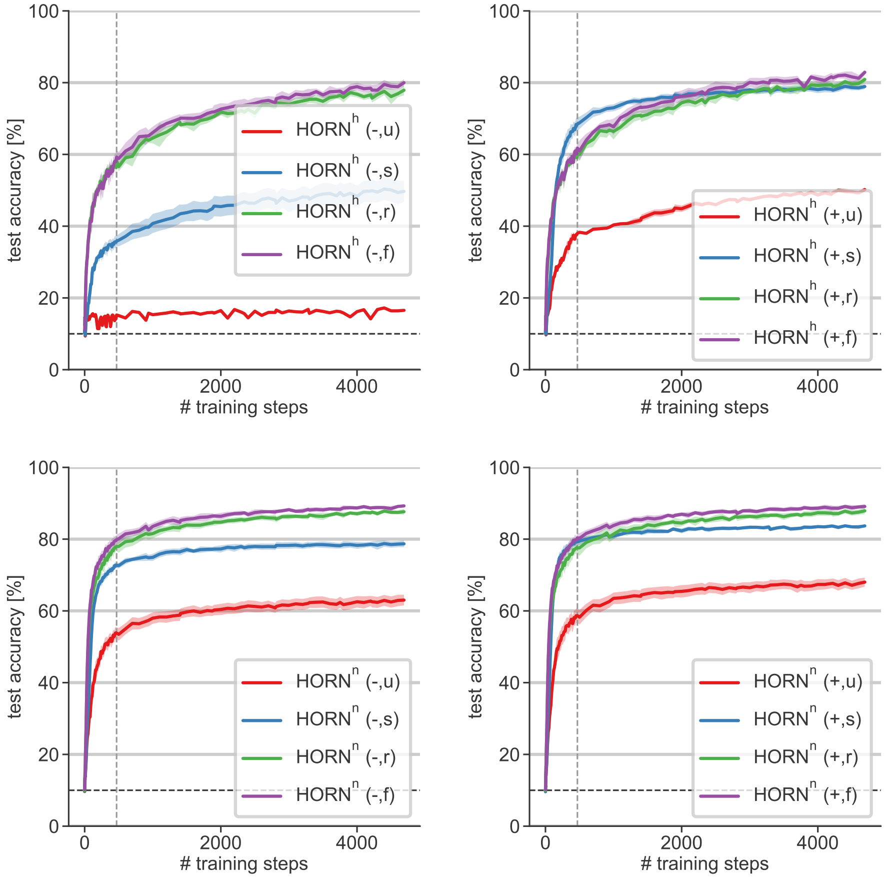


**Fig. S17**. Learning dynamics of HORN networks (32 nodes) as in Fig. S16 trained on psMNIST for 10 epochs.

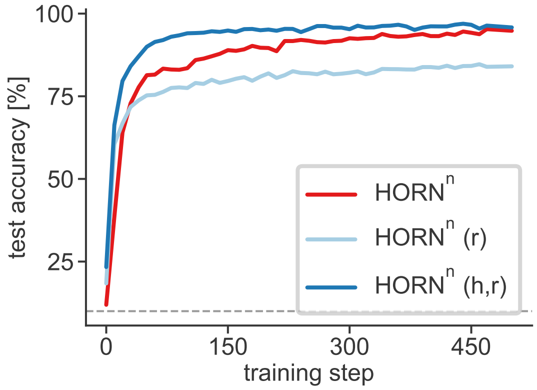


**Fig. S18**. Learning dynamics of HORN$^n$ network (93 nodes) when trained on sMNIST from scratch (red line), or with pre-training on LSDSb data set under different training conditions: (i) Only readout weights trained (light blue), and (ii) readout and recurrent weights trained (dark blue).

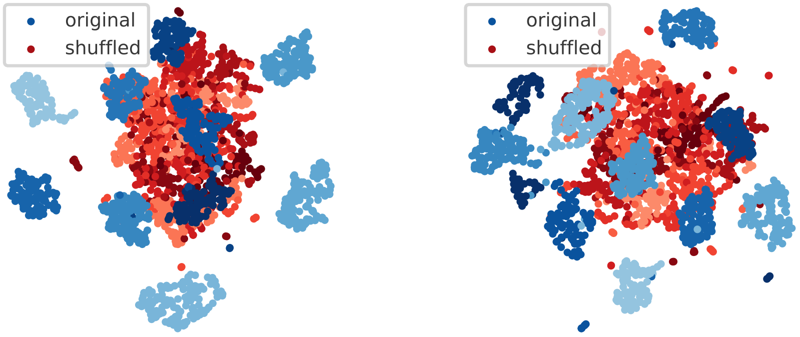


**Fig. S19**. UMAP projection of homogeneous (left, 93 nodes) and heterogeneous HORN (right, 93 nodes) network state at the roadout time ($t = 784$) after training on sMNIST for 10 epochs. Activity elicited by sMNIST stimuli shown in blue shades, acitivity elicited by psMNIST stimuli shown in red shades. MNIST digit class coded in color lightness in both cases.

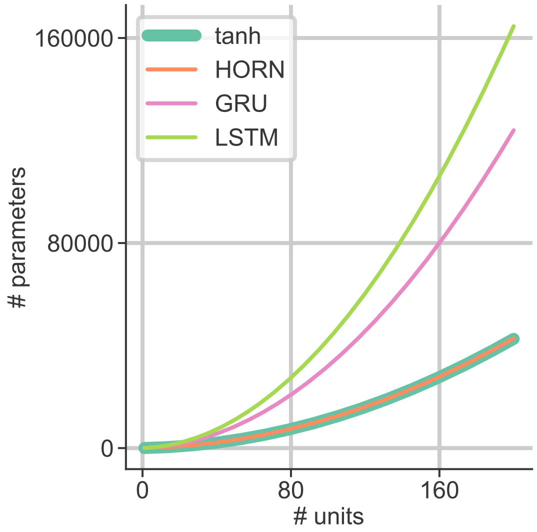


**Fig. S20**. Number of trainable system parameters as a function of network size for different RNN architectures and sMNIST or psMNIST classification (input signal dimensionality 1, readout signal dimensionality 10).

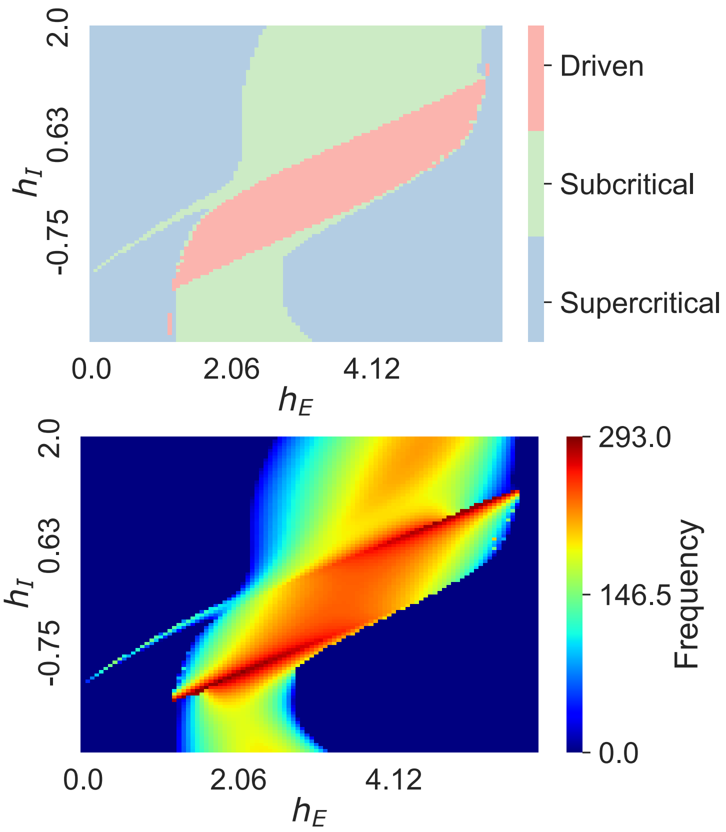


**Fig. S21**. Wilson-Cowan dynamics in $h_E$-$h_I$ subspace. Top: Bifurcation diagram, blue: supercritically damped harmonic oscillations, green: subcritically damped harmonic oscillations, red: driven subcritically damped harmonic oscillations. Bottom: Oscillation frequencies (frequency units arbitrary).

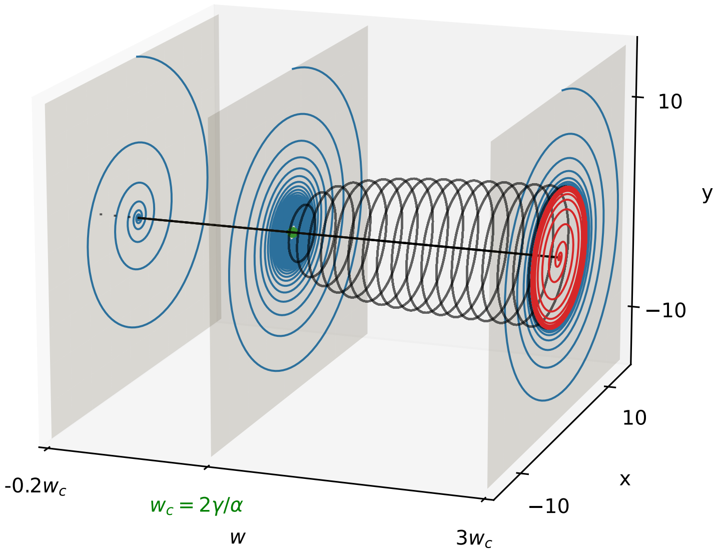


**Fig. S22**. Hopf bifurcation of a DHO node ($\omega = 1$, $\alpha = 1$ and $\gamma = 0.1$) with velocity feedback term $w$ at a critical value $w_c = 2\gamma/\alpha = 0.2$. Blue lines: node relaxation dynamics from the initial condition $(x_0, y_0) = (0, 15)$ for different values of $w$. Green dot: Hopf bifurcation point at $w = w_c$. Black circles: limit cycle solutions with a radius depending on $w$. Red line: Relaxation dynamics from the initial condition $(x_0, y_0) = (0, 0.1)$ for $w = 0.6$. Black line: $(x, y) = (0, 0)$.

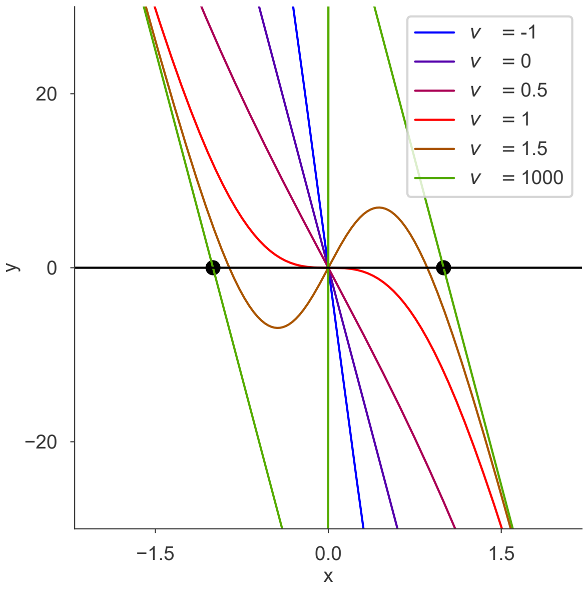


**Fig. S23**. Intersections between the nullclines of DHO system dynamics. Black curve: $\dot{x} = 0$ (S10). Other curves: $\dot{y} = 0$ (S9, $\alpha = \omega = 1$) for different values of amplitude feedback parameter $v$ (color coded). Black dots marks the points $(\pm\frac{\alpha}{\omega^2}, 0)$.

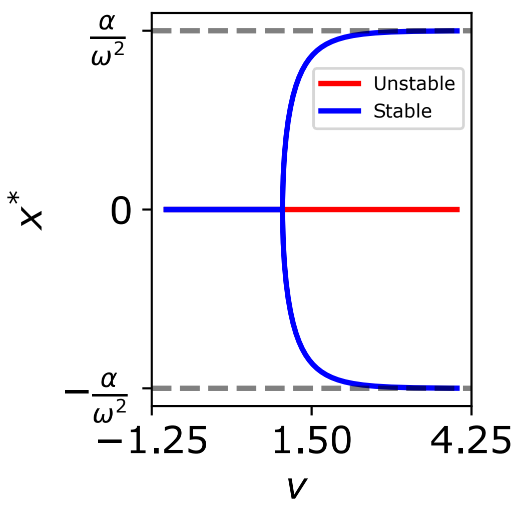


**Fig. S24**. Bifurcation diagram of DHO node ($\alpha = \omega^2 = 1$) dynamics under changes in the amplitude feedback parameter $v$ in $(v, x^*)$-subspace. Pitchfork bifurcation point at $v_c = \frac{\omega^2}{\alpha}$.

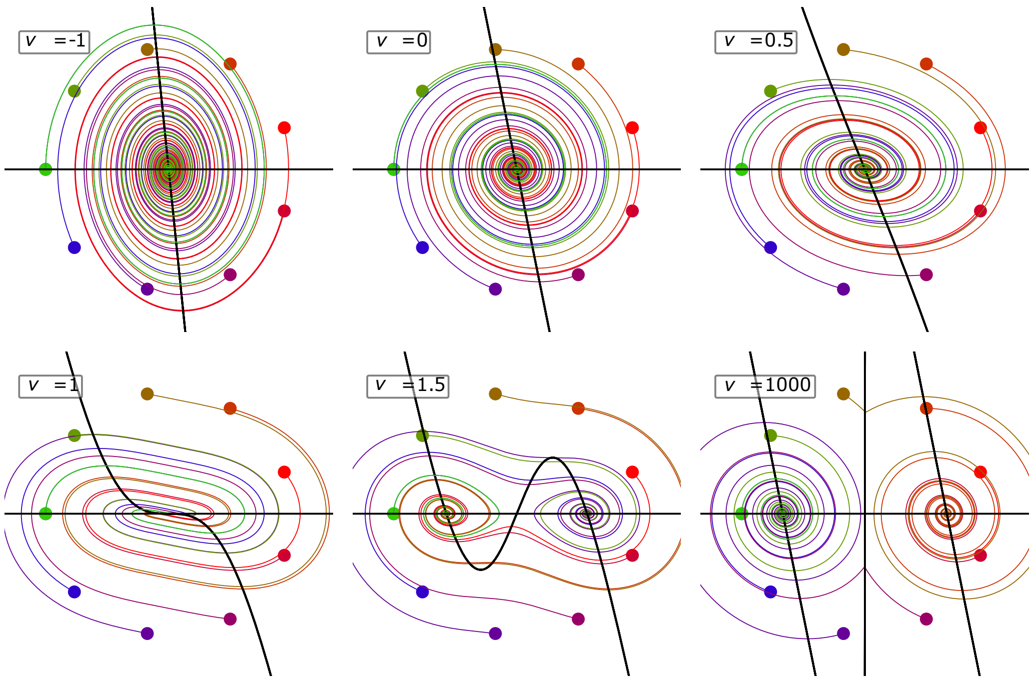


**Fig. S25**. Phase portraits of relaxation dynamics of DHO node ($\omega = 1$, $\alpha = 1$, $\gamma = 0.1$) for different values of the amplitude feedback parameter $v$. Different initial conditions color coded.

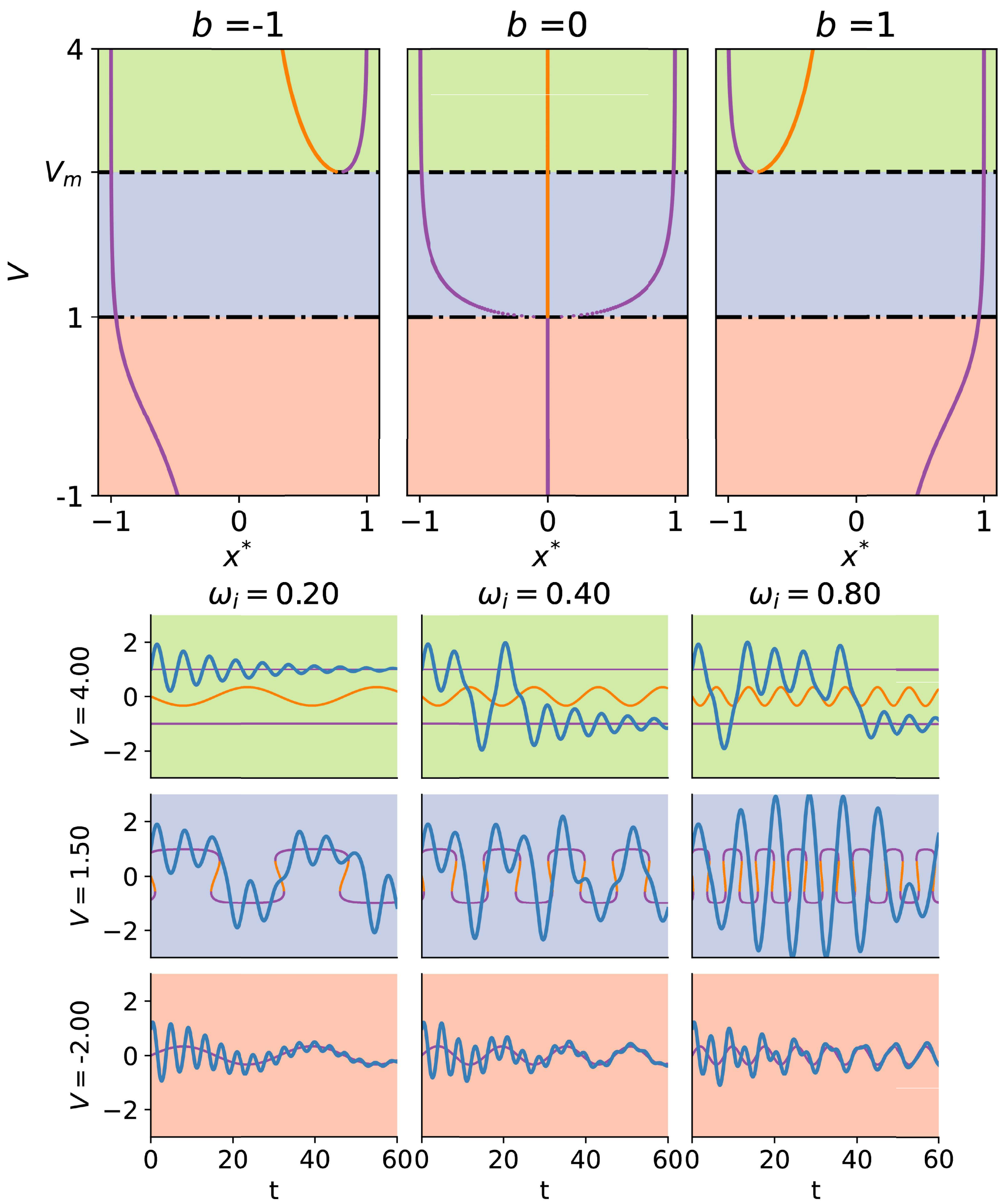


**Fig. S26**. Symmetry breaking of the pitchfork bifurcation given a constant external input and dynamics of nodes subjected to harmonic inputs. Top: Bifurcation diagrams of DHO nodes for different values of constant external input $b$, where $\omega = 1, \alpha = 1 and \gamma = 0.05$. Dashed line: Local minima $v = v_m$ (S16). Dash-dotted line: Pitchfork bifurcation point at $v = v_c = 1$. Orange curve: unstable branch, purple curves: stable branches. Colored regions indicate qualitatively different configurations o moving fixed points, due to harmonic input. Green: 3 fixed points, blue: 3 to 1 fixed points. Red: 1 fixed point. Bottom: DHO moving fixed points and an example of a node trajectory for harmonic input with frequency $\omega_i$. Purple: stable branch, orange: unstable branch, blue: node trajectory. Background colors indicate corresponding region on the top panel.

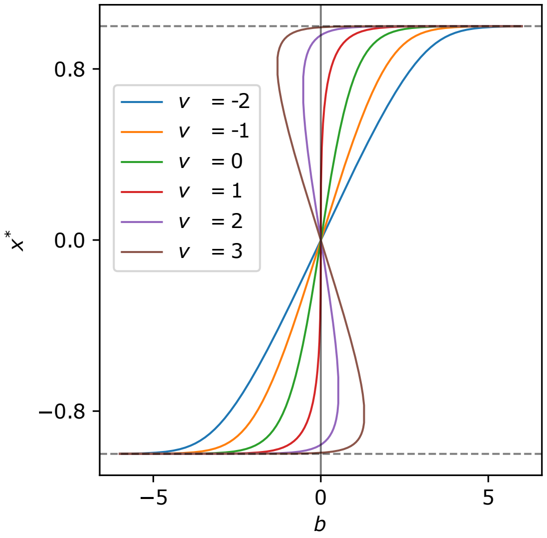


**Fig. S27**. Position of DHO nodes ($\omega = 1$, $\alpha = 1$, $\gamma = 0.01$) fixed points for a constant input of amplitude $b$ and different values of the amplitude feedback parameter $v$ (color coded).

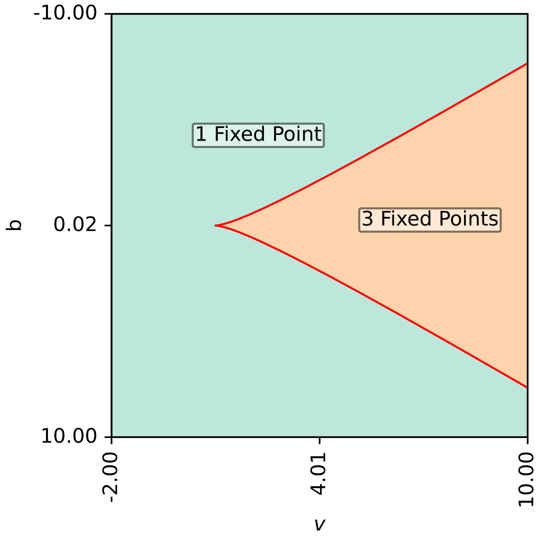


**Fig. S28**. Cusp bifurcation of the dynamics of a DHO node ($\omega = 1$, $\alpha = 1$, $\gamma = 0.01$) subject to an external constant input of amplitude $b$ with amplitude feedback strength $v$, shown in $(v, b)$-space.

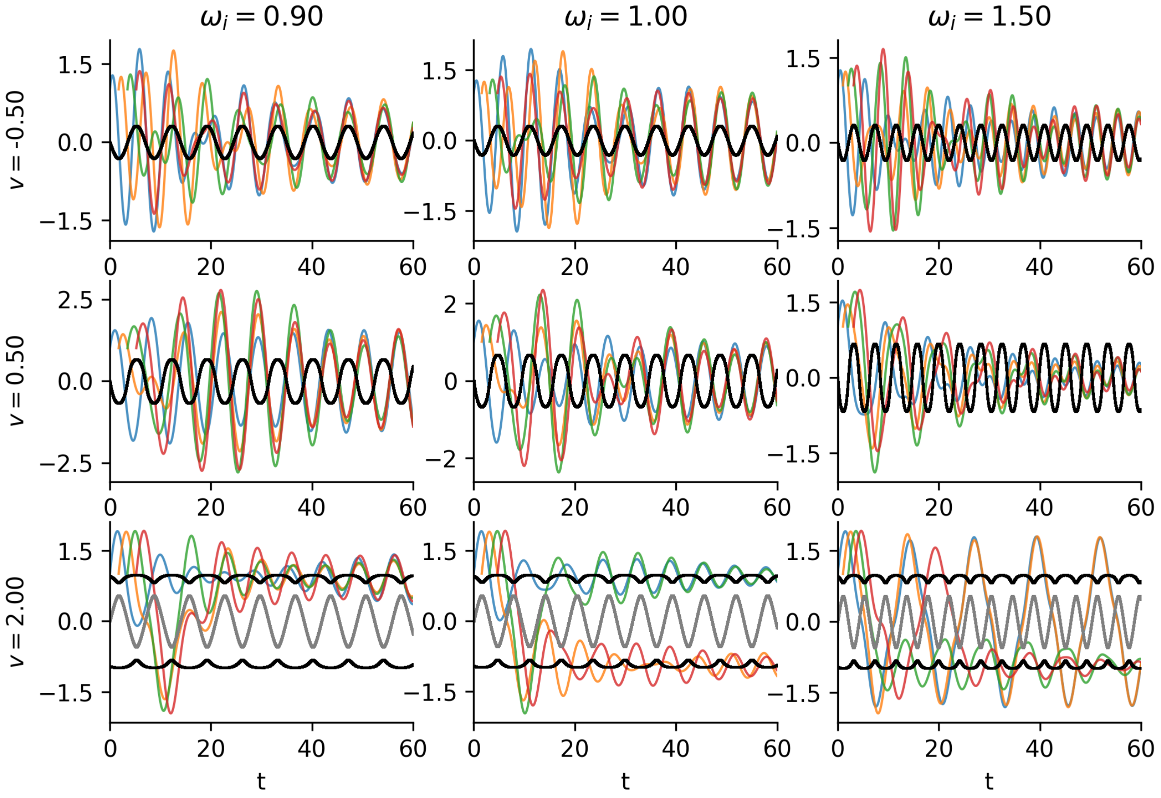


**Fig. S29**. DHO node ($\omega = 1$, $\gamma = 0.05$, $\alpha = 1$) relaxation dynamics dependence on the values of initial time $t_0$. Panels show cases for different values of harmonic input frequency $\omega_i$ and amplitude feedback values $v$. Blue: $t_0 = 0$, orange: $t_0 = \frac{\pi}{2\omega_i}$, green: $t_0 = \frac{\pi}{\omega_i}$, red: $t_0 = \frac{3\pi}{2\omega_i}$, black: stable moving fixed point, gray: unstable moving fixed point.

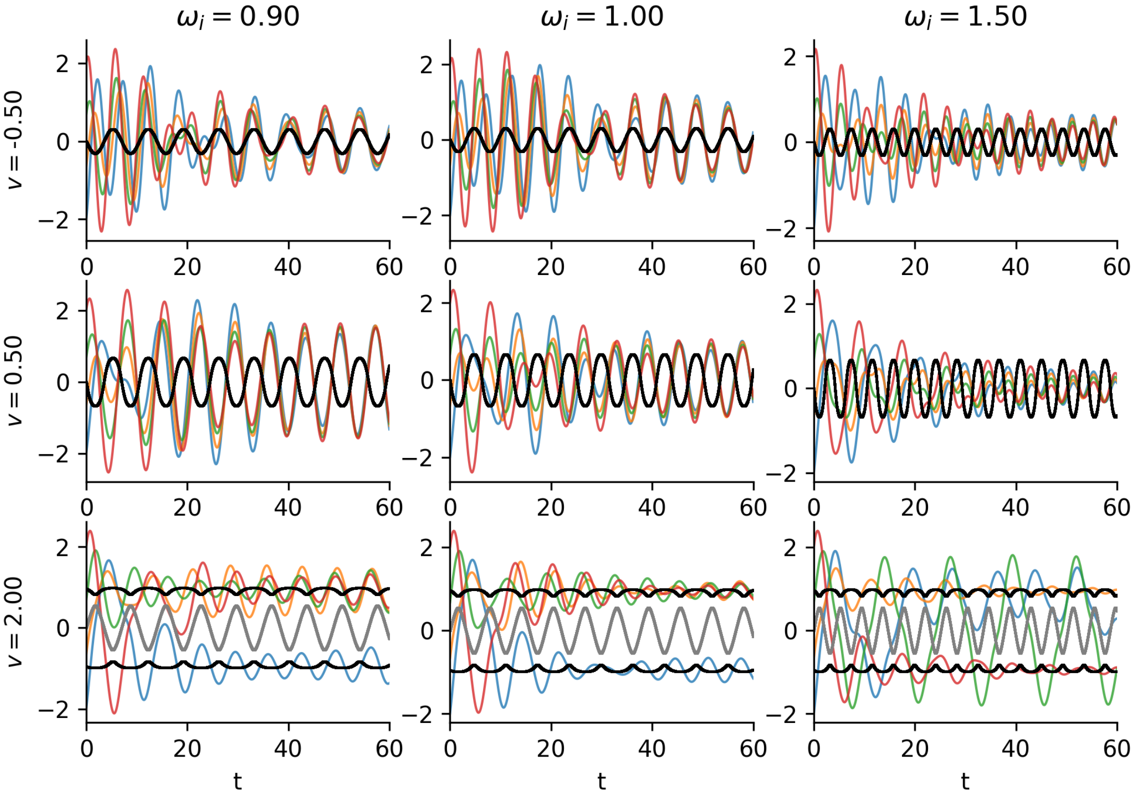


**Fig. S30**. DHO node ($\omega = 1$,$\gamma = 0.05$, $\alpha = 1$) relaxation dynamics dependence on the values of initial amplitude $x_0$. Panels show cases for different values of harmonic input frequency $\omega_i$ and amplitude feedback values $v$. Blue: $x_0 = -2$, orange: $x_0 = -1$, green: $x_0 = 1$, red: $x_0 = 2$, black: stable moving fixed point, gray: unstable moving fixed point.

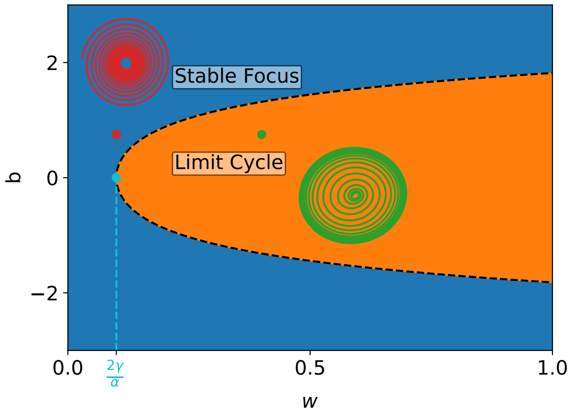


**Fig. S31**. Hopf bifurcation of the dynamics of a DHO node ($\omega = 1$, $\alpha = 1$, $\gamma = 0.05$) in the $(w, b)$ parameter space, where $w$ denotes velocity-feedback and $b$ constant input amplitude. Colored lines show example trajectories on the phase space for each region.

Table 1: Parameter values for studied RNN architectures on different data sets. Columns $\omega_p$, $\gamma$, $\alpha$ show parameter values for HORN networks. First value in each column specifies parameter value for homogeneous HORN, values in parentheses denote value ranges for heterogeneous HORN. Parameter values of the natural frequency are given in period units of the input, and $\omega = 2\pi/\omega_p$. Columns $h$, $a$ show parameters for leak RNN.

| **Dataset** | $\omega_p$ | $\gamma$ | $\alpha$ | $h$ | $a$ |
|---|---|---|---|---|---|
| sMNIST | 28 ([28, 85]) | $10^{-2}$ ([$10^{-4}$, $2 \cdot 10^{-2}$]) | $4 \cdot 10^{-2}$ ([$10^{-2}$, $1.6 \cdot 10^{-1}$]) | 0.2 | 0.2 |
| psMNIST | 7 ([6, 22]) | $5 \cdot 10^{-2}$ ([$1.5 \cdot 10^{-2}$, $9 \cdot 10^{-2}$]) | $2.5 \cdot 10^{-1}$ ([$9 \cdot 10^{-2}$, $3.6 \cdot 10^{-1}$]) | 0.5 | 0.3 |
| EMNIST | 28 ([30, 85]) | $5 \cdot 10^{-2}$ ([$10^{-3}$, $5 \cdot 10^{-2}$]) | $10^{-2}$ ([$2.5 \cdot 10^{-2}$, $4 \cdot 10^{-2}$]) | 0.1 | 0.2 |
| LSDS | 28 ([28, 85]) | $10^{-2}$ ([$10^{-4}$, $2 \cdot 10^{-2}$]) | $4 \cdot 10^{-2}$ ([$10^{-2}$, $1.6 \cdot 10^{-1}$]) | 0.2 | 0.2 |
| SDDS | 63 ([42, 126]) | $5 \cdot 10^{-4}$ ([$5 \cdot 10^{-5}$, $1.5 \cdot 10^{-3}$]) | $10^{-2}$ ([$2.5 \cdot 10^{-3}$, $2.5 \cdot 10^{-2}$]) | 0.1 | 0.2 |
| MNIST | 18 | $10^{-3}$ | $10^{-2}$ | – | – |

**Movie S1.** Temporal evolution of the activity of the nodes of a 196 node HORN$^h$ trained on geometrically organized MNIST input for 10 example digits. Nodes are arranged on a $14 \times 14$ matrix according to the geometric location of the nodes' $2 \times 2$ receptive fields in the $28 \times 28$ stimulus matrix. Each trial lasts 150 time steps, and the stimulus is presented during the first time step and then switched off. Node amplitudes are color-coded (color scale $[-0.1, 1.0]$ for first 50 time steps and $[-2.0, 2.0]$ for the remaining time steps to enhance visibility). Current time step $t$ and stimulus identity shown at the top, asterisk marks stimulus presentation.

**Movie S2.** Temporal evolution of the phases of the nodes of a 196 node HORN$^h$ trained on geometrically organized MNIST input for 10 example digits. Node phases $\phi \in [-\pi, \pi]$ are color-coded. Trials and node placement correspond to the ones in Movie S1.
